# CAcnesDB: A database of *Cutibacterium acnes* with Integrated Functional Insights derived from Multi-modal Genome Annotation

**DOI:** 10.1101/2025.11.20.689451

**Authors:** Bandana Pandia, Dheemanth Regati, Nihesh Rathod, Santhosh Sankar, Sneha Vasudevan, R Sowdhamini, Nagasuma Chandra

**Affiliations:** Department of Biochemistry, Indian Institute of Science, Bangalore, Karnataka, 560012, India; IISc Mathematics Initiative, Indian Institute of Science, Bengaluru, Karnataka, 560012, India; Centre for Biosystems Science and Engineering, Indian Institute of Science, Bengaluru, Karnataka, 560012, India; National Centre for Biological Sciences, TIFR, GKVK Campus, Bellary Road, Bangalore 560065, India; Department of Electrical Communication Engineering, Indian Institute of Science, Bengaluru, Karnataka, 560012, India

## Abstract

*Cutibacterium acnes* is a predominant member of the human skin microbiome, generally as a beneficial or a benign inhabitant, which under specific conditions transforms into a pathogen, causing acne and associated inflammatory responses. While there is a wealth of data on the microbiology and genomics of the microbe, there is a lack of comprehensive annotation of the proteins coded by the genome, slowing progress in understanding its unique metabolic versatility and survival in host hypoxic sebaceous niches. We configured an integrated pipeline for functional annotation of the *C. acnes* genome, combining gene calling, domain and ontology mapping, KEGG pathways assignment, proteome-scale structural modeling, predicting Pockets and associating the small-molecule metabolites based on binding site similarity to facilitate structure-to-function annotation. Structural predictions using ColabFold covered nearly the full proteome, with 67.4% of models showing high confidence. To infer protein function, ligand-binding sites in the *C. acnes* modeled proteome were predicted using a consensus of three independent algorithms, PocketDepth, SiteHound, and FPocket. The resulting pockets were matched with known small-molecule binding sites in PDB to map potential ligands, resulting in the *C. acnes* predicted metabolome (CApM) comprising 1,954 ligands across 1,865 proteins. CAcnesDB is a comprehensive up-to-date database of the *C. acnes* genome, hosting functional annotation from multiple levels of input and analyses, covering 2297 protein-coding genes.

**Impact Statement:** *Cutibacterium acnes* (*C. acnes*) is implicated in acne vulgaris, a common skin condition affecting millions of people worldwide, yet the absence of a unified genomic resource limits comprehensive understanding of the underlying biology. In this work, we introduce CAcnesDB, a genome-wide annotation resource built through multi-modal integration of sequence- and structure-based analyses, structural proteome models, predicted binding sites associated with small-molecule metabolites, functional classes, protein–protein interactions, and pathways. This resource consolidates enriched annotations to enable systematic exploration of *C. acnes* biology and pathogenic mechanisms. We showcase its utility through detailed analyses of membrane transporters and hypothetical proteins, highlighting novel functional insights into this skin microbe. Given the clinical relevance of *C. acnes* in skin health and its role in skincare product development, CAcnesDB represents a valuable platform for both basic research and translational applications.

## Introduction

*Cutibacterium acnes* is a Gram-positive, facultative anaerobic bacterium that resides near the sebaceous glands inside the pilosebaceous units (PSU) present on the skin^1,2^. It has saprophytic properties and lipolytic activity, owing to which it can acquire nutrients from the secretions of sebaceous follicles, such as sebum lipids. It is a major human skin commensal and produces short-chain fatty acids (SCFAs) such as propionic acid that maintain the skin’s acidic pH and inhibit the growth of pathogenic microbes. The latest classification of *C. acnes* strains includes three subspecies: *C. acnes acnes* (phylotype I, ribotype (RT)-RT1, RT3, RT4, RT5, RT8, RT16, RT532), *C. acnes defendans* (phylotype II, ribotype - RT2, RT6), and *C. acnes elongatum* (phylotype III, ribotype-RT9)^3^. Phylotypes II and III are often found on healthy skin and are non-acneic strains^4^. In contrast, subtypes IA-1 (RT1, RT5, RT532) and IA-2 (RT1, RT4, RT5) are acneic strains and are associated with enhanced virulence, inflammatory potential, and antibiotic resistance^3,5^. The first report of the genome sequence annotation of *C. acnes* was in 2004 (Bruggemann et al. 2004; KPA171202 strain 2.56 Mbp, 60% GC content, phylotype IB and ribotype 1)^6^. Since then, about 190 genome sequences of various strains (complete and draft) have been determined, which have indicated the core genome of *C. acnes* that includes both host-beneficial and host-detrimental genes^1,4^. Although this microbe is a part of the healthy skin flora, in an opportune situation, it turns into a pathogen, causing proinflammatory skin diseases, mainly acne vulgaris and implant-associated infections^1^. A comparative genome study between strain 266 of phylotype IA and strain KPA171202 of type IB, identified a number of island-like genomic regions encoding a variety of putative virulence-associated factors, such as dermatan-sulfate adhesin, polyunsaturated fatty acid isomerase, iron acquisition protein HtaA, and lipase GehA, which were upregulated in strain 266, as well as genes involved in various transport systems and metabolic pathways^7^.

Comprehensive genome annotation is a key requirement for understanding the capability of the organism from a wide range of perspectives, including metabolic adaptation to physiological stimuli, host environmental niches and abiotic stresses, as also for genome comparisons and host-pathogen interactions and biomedical applications, including diagnosis, drug and vaccine discovery. For several organisms, rich annotation using bioinformatics methods has led to the development of databases such as PANTHER, IMG, KEGG, and organism-specific resources such as FlyBase, WormBase, Mycobrowser, TBDB, EcoCyc, and ECMDB, which have greatly aided in collating and organizing information for targeted organisms^8,9^. Although *C. acnes* has been a microbe of interest given how widespread the acne condition is, there is no resource providing comprehensive genome annotation. In the present study, we address this gap and have carried out multimodal multilevel bioinformatics analysis to comprehensively characterize the whole genome and present the results in a dedicated database, CAcnesDB. We anticipate that CAcnesDB will benefit researchers in gaining a more holistic understanding of the diversifying role of *C. acnes*, aiding in the experimental design to accelerate the identification of drug-targets in *C. acnes*.

## Methods

### Sequence Analysis, Gene Finding, Gene Mapping, and Annotation Categorization

The whole genome of *C. acnes* KPA171202 was subjected to systematic identification, characterization, and confirmation for each gene candidate. Several tools and algorithms, such as Prokka^10^, PGAP^11^, Pannzer2^12^, InterProScan^13^, BlastP^14^, HHblits^15^, KEGG tools^16^, STRING database^17^, VirulentPred^18^, P2RP (Predicted Prokaryotic Regulatory Proteins)^19^, and DeepTFactor^20^ were included in the sequence-to-function assignment protocols (Supplementary Methods Doc1). The full list of tools, algorithms, and databases used for this study is listed in Supplementary Sheet S1. Prokka (Rapid Prokaryotic Genome Annotation) and PGAP (Prokaryotic Genome Annotation Pipeline) were used to generate a first draft of the revised annotation for the whole genome. Both PROKKA and PGAP employ the BLAST (Basic Local Alignment Search Tool) algorithm to search against broader databases of proteins, i.e., UniProt^21^ and RefSeq^22^, respectively. For clarity and mapping consistency, the gene entries in the KEGG database^23^ were used as the reference. That resulted in a total of 2369 genes, out of which 55 (6 rRNA + 47 tRNA + 2 tmRNA) were non-coding genes, 17 were pseudogenes, and 2297 genes were protein-coding genes (CDS). All the CDS were further categorized into three subgroups: ‘hypothetical’, ‘putative,’ and ‘known’ proteins based on the search results of the matching words from the existing ‘GenBank’ definition, as illustrated in Figure S1(a). The gene grouping was done following the annotation guidelines for prokaryotes, as reported by the NCBI, https://www.ncbi.nlm.nih.gov/genbank/genomesubmit_annotation/. Proteins with ‘putative’ and ‘hypothetical’ terms in their GenBank definition were categorized into the ‘hypothetical’ subgroup. Further, known proteins having the ‘conserved’ word were categorized into the ‘putative’ subgroup. The STRING database was utilized to add the protein–protein interactions, functional annotations, and corresponding KEGG pathways of the *C. acnes* KPA171202 proteome, https://version-12-0.string-db.org/organism/STRG0A28GPG. After the automated genome annotation, results were corroborated with pre-existing GenBank definitions for each gene, where available.

### Genome Annotation From Structures

Various Software tools, algorithms, and databases used for the ‘annotation from structures’ study, such as ColabFold (MMSeq2 + Alphafold2)^24^, TM-align^25^, PocketDepth^26^, FPocket^27^, SiteHound^28^, SCOP2^29^, and FLAPP^30^, are listed in Supplementary Sheet S1. Briefly, the pipeline consists of (a) prediction of protein structure, (b) prediction of pockets, and (c) comparison of binding sites for individual protein sequences in *C. acnes* KPA171202, as shown in Figure S2.

### Structural Models

The genome of the *C. acnes* KPA171202 encodes 2297 protein-coding genes. Structural models were generated using ColabFold, based on the fast homology search of MMseqs2 for generating Multiple Sequence Alignments (MSAs) of the query protein, which is taken as input for the second algorithm, Alphafold2, to generate predicted structures, from which the best-ranked model is selected for each query protein. The scan ColabFold predicted models for 2280 proteins, nearly the whole proteome (excluding 17, of which 15 proteins have a polypeptide length greater than a thousand residues).

### Quality of the Predicted Structures

In a modeled protein structure, each amino acid is associated with a per-residue Local Distance Difference Test (pLDDT) score. The pLDDT values for each residue were utilized to assess the quality of the modeled *C. acnes* proteome with the reported thresholds, and based on the confidence criteria suggested by AlphaFold developers (highly accurate: regions with pLDDT scores ≥ 90; good confidence: 90 > pLDDT score ≥ 70, low confidence: 70 > pLDDT ≥ 50; uninterpretable: pLDDT < 50), and retained only those regions with pLDDT > 50. Structural alignments were performed using TM-align against non-redundant protein structures present in PDB (m=26,775), to assign SCOP2-fold, family, and superfamily information to *C. acnes*’ modeled proteome (n=2280), which is detailed in Supplementary Methods Doc1.

### Pocketome detection

Consensus small molecule binding sites (*C. acnes* pocketome) were predicted using three methods: PocketDepth (Geometry, Centrality-based), FPocket (Voronoi tesselation-based), and SiteHound (Energy-based). PocketDepth is an in-house method that identifies a probable binding pocket in a given structural model by depth-based clustering sub-spaces in possible pocket candidates and selecting those with highly central sub-spaces to define putative pockets^26^. SiteHound identifies a putative pocket by finding the regions on the protein surface that are energetically favorable using Molecular Interaction Fields (MIF)^28^. FPocket depends on Voronoi tessellation, constructs a set of small alpha spheres in proteins that connect four atoms so as to make them equidistant to the central alpha sphere^27^. These three independent methods that run on different principles were used to detect the pockets. Putative small-molecule binding sites, which were predicted by all three, were considered to increase the probability of their existence. Predicted pockets in which at least 70% of residues had a pLDDT score ≥ 70 were designated as consensus pockets (CPs) and retained for downstream analysis.

### Binding site comparison

The predicted binding sites (pockets) are aligned against non-redundant PDB binding sites (NrSites) using FLAPP, an in-house fast atomic-level alignment algorithm^30^. FLAPP compares two binding sites based on their size and residue properties and outputs three values: alignment length, F_min_, and F_max_. In addition, it also reports actual residue-residue correspondences, where alignment length is the number of residues common between two sites. F_min_ and F_max_ is a score representing the structural similarity between the sites with respect to the smaller and the larger sites respectively. F_min_ ≥ 0.4 and N ≥ 6 has been reported as suitable cutoffs^30^ to indicate that the two binding sites are similar, and hence, the same cutoff was used throughout the analysis.

### Structure-to-Function Association

The individual steps involved in associating a function based on the structures are illustrated with an example. Similar analyses and annotations have been derived for the rest of the *C. acnes* proteome. PPA0018 was identified to be a ribokinase (RbsK), having a length of 309 amino acid residues (AA). The top-ranked predicted structural model covered the entire polypeptide with a pLLDT ≥ 70 for 302 AA amounting to 97.7% of ‘high confidence’ coverage. Seven Consensus Pockets (CPs) were identified from the consensus pocket-detection pipeline (Pocket-0, 1, 2, 3, 6, 7, and 10). The closest FLAPP hit for pocket-1 in PDB was identified to be ‘4XCK_RIB_A_401.pdb’, where ‘RIB’ stands for ‘alpha-D-ribofuranose’, based on which a functional association of ligand ‘RIB’ was made for ‘Ribokinase’. Similarly, functional descriptions were assigned or confirmed for each of the associated ligands with their corresponding proteins. The ligands identified for the entire proteome, together, constitute the small-molecule binding metabolites or ligands, thus called *C. acnes*’ predicted Metabolome (CApM). Autodock^31^ was used to calculate the binding energy (free energy change, ΔG), to assess the binding affinity of a mapped ligand to the predicted pocket of a modeled protein.

### Assessment of the annotated functional description

Iterative manual curation and literature-based up-to-date annotation were carried out to corroborate the assigned annotation across all the evidences for each individual protein. Additionally, an evidence scoring schema was devised to score the reliability of the assigned functional annotation (Supplementary Methods Doc1).

### Database and Full-Stack Web Development

The CAcnesDB database was developed using Node.js, Express.js, JavaScript, CSS, and HTML to create a data-centric web application (listed in Sheet S1). The web interface is developed using Node.js (https://nodejs.org/en). Node.js^32^ is a server-side JavaScript runtime environment that builds scalable and efficient applications that interact with databases and handle data operations. Express.js^33^ is a web application framework for Node.js that simplifies the process of building robust and modular web applications. JavaScript was used for both server-side (Node.js) and user-side (browser) scripting. The web application used CSS (Cascading Style Sheets) to style visual aspects such as layout, colors, and typography. HTML (Hypertext Markup Language) was used to structure and present the contents on the web. The combination of these technologies was utilized to create a Full-Stack Web Development, which facilitates a seamless user experience, managing and storing data locally in JSON files as well. Additionally, STRING API^17^ was integrated with the CAcnesDB, which provides an interactive display for protein-protein network visualization. Mol*(https://molstar.org/)^34^ was used to represent the protein structures in 3 dimensions. It is a collection of tools that have been designed to visualize and analyze molecular structures in 3D space. It provides interactive and dynamic representations of the molecular structures. The modeled structures and sequences can be downloaded in PDB and FASTA formats, respectively. The website allows a user to sort the tables and search for information. The database can be queried using PPA IDs, gene names, UniProt IDs, Pfam IDs, RefSeq IDs, and CAcnesDB functional classification. Database URL: https://proline.biochem.iisc.ac.in/CAcnesDB/

## Results

### Overview of the genome annotation

We configured a computational pipeline for genome annotation, comprising sequence analysis of the gene products, protein structure analysis, binding site structural analysis, site structure-based ligand identification, and database development (Figure 1). The assignment of functional descriptions to each protein was systematically made, based on the results obtained from individual analyses. The annotations were manually cross-checked with the existing ‘GenBank’ definitions obtained from the entries in KEGG, and updated with new information from the various steps comprising the protein characterizations based on sequences and modeled structures. Next, we consolidated the evidence for functional annotation from multiple levels of analyses, including from a structure-based ligand prediction step as well as consolidation of findings in literature where available, to get updated genome characterization and protein annotation at high resolution. We indicate the confidence from the multiple evidences for the functional descriptions, using an annotation scoring schema. We present a database, CAcnesDB, to describe the functional annotation. We showcase the usefulness of the database through selected examples belonging to the categories of ABC transporters and hypothetical proteins.

**Figure 1:**
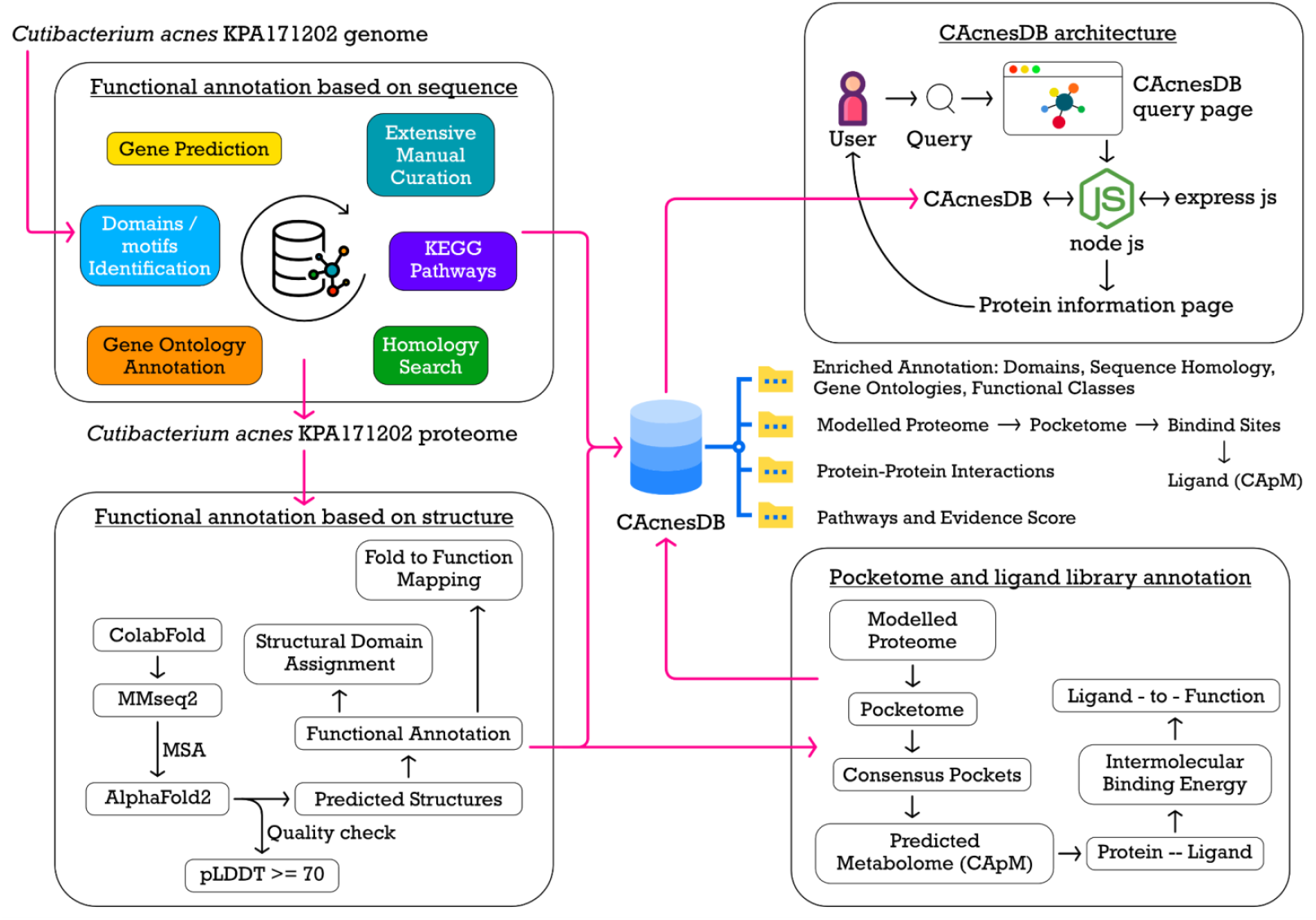
A schema representing an integrated pipeline comprising tools and databases utilized for genome annotation from sequences and structures, and construction of the CAcnesDB. Different aspects of annotation are illustrated in the schema.

### Summary Statistics

The multi-step annotation pipeline for annotation based on sequences included gene calling (with PROKKA and PGAP), and an integrated usage of various tools and databases to confirm a revised annotation. This included identification of domains, Gene Ontologies, functional categories from descriptions, KEGG orthology mapping, and assigning to the KEGG pathways. The detailed report on sequence-to-function results is provided in Supplementary Results Doc2. In brief, of the 2297 protein coding genes in the genome, domains were assigned for 1958, sequence-sequence homology mapping was obtained for 1721, profile-profile sequence matches obtained for 2297 and structural models were predicted for 2280, binding sites predicted for 2205, ligands predicted for 1865, protein-protein interactions predicted for 1890 proteins, put together providing functional annotation for 2072 proteins (Detailed statistics in Sheet S1-summary stats). Overall, the CAcnesDB database contains annotations for the *C. acnes* KPA171202 strain, associated KEGG orthologues, KEGG pathways, GO terms (BP, MF, and CC), SCOP2 hierarchies, functional class, and evidence-based annotation scores. The general statistics of the results in CAcnesDB are listed in Table 1. A comparison of the results in CAcnesDB with the corresponding data from public repositories is provided in Supplementary Table S1.

**Table 1:** Summary statistics of the availability of results from each analysis carried out in the annotation of the *C. acnes* KPA171202.

|  | **CAcnesDB** |
| --- | --- |
| **# Protein coding genes (CDS)** | 2297 |
| **# Pseudogenes** | 17 |
| **# Protein non-coding genes** | 55 (6 rRNA + 47 tRNA + 2 tmRNA) |
| **# Protein with definite functional assignments (absence of ' Putative/ Probable/ Predicted/ Hypothetical')** | 1826 |
| **# Protein with 'Probable/ Putative/ Predicted' functional assignments** | 215 (194 'putative' + 14 'probable' + 7 'predicted') |
| **# Hypothetical/ Uncharacterized/ Domain of Unknown (DUF) proteins** | 256 (225 are newly annotated) |
| **# Protein with gene symbols** | 823 |
| **# Protein with assigned domains** | 2085 (InterPro, Pfam, Gene3D, CDD, Coils, HAMAP, PRINTS, PrositePatterns, PrositeProfiles, SFLD, SMART, superfamily, Tigrfam, MobiDB, Panther) |
| **# Protein with assigned Gene Ontology (GO) (BP or MF or CC)** | 1686 (BP: 1036, MF: 1566, CC: 326) |
| **# Protein with assigned KEGG orthology (KO)** | 1503 |
| **# Protein with assigned KEGG pathways/modules** | 1596 (KEGG pathways + KEGG modules) |
| **# Structural models for proteins** | 2280 (predicted models from ColabFold) |
| **# Protein with SCOP domains** | 1205 (Superfamily and/or FOLD and/or Family) |
| **# Protein with predicted ligands** | 1865 |
| **# Proteins with predicted ligands with intermolecular binding energy $\leq -5$ kcal/mol** | 1562 |
| **Availability of Functional for the whole genome of *C. acnes*** | Yes |
| **# Functional Classes** | 9 Functional Classes |
| **# Proteins with Functional Classes** | 2273 |
| **Availability of Annotation-confidence schema** | Yes |

### Sequence-based Annotations

The sequence-based function annotation for a gene comprised information from one or more of the following: (a) Gene finding: functional annotation obtained from the Prokka and PGAP analyses, (b) Homology search: functional description of the top-hits obtained sequence-sequence and profile-profile alignments, (c) Gene Ontology (GO): obtaining functional role from the combined knowledge of Biological Processes (BP), Molecular Function (MF), and Cellular component (CC), (d) Pathway analysis: deriving the functional role from KEGG orthology (KO) and pathway information, (e) Domains/Signatures/Motifs: Establishing the functional capacity from the combined domains, motifs and signatures associated with a protein sequence, (f) Virulence: identifying the functional capacity of a protein for its virulence or regulatory activity, and (g) Miscellaneous: Literature-based annotation derivation for other available protein functional terms.

Sequence-based homology identified functions for 1,721 CDS, while profile-profile searches yielded functional descriptions for 2,297 genes. Domains/ Motifs were be identified for a major portion of the *C. acnes* genome. Out of 2297 protein-coding genes (CDS), 1872 (≈ 81.5% coverage) were associated with Pfam domain IDs, and 938 (≈ 40.8% coverage) were associated with CDD domain IDs. The overall output from the InterProScan is listed in Sheet S1 (Summary stats).

Figure S3(a) summarizes the distribution of three GO terms across the whole genome in *C. acnes* KPA171202. In total, 149 genes could be associated with all three GO terms. Figure S3(b) displays the distribution of GO (BP) categories among the 1,036 annotated predicted protein-coding genes. The most abundant GO:BP were transmembrane transport, carbohydrate metabolic process, and cellular amino acid metabolic process. Figure S3(c) displays the distribution of GO (MF) categories for 1,566 annotated predicted protein-coding genes. The top GO: MF terms were transferase activity, hydrolase activity, and oxidoreductase activity. Figure S3(d) displays the distribution of GO (CC) categories for 326 protein-coding genes. The most abundant GO: CC were the plasma membrane and ribosome, followed by the organelle and the extracellular region.

Proteins’ function culminates from the interactions between proteins, either in direct physical complexes or indirect associations, available in STRING-db, a database that contains protein-protein interaction networks for >10,000 diverse genomes^17^. The coordinated functions performed by a group of proteins that facilitate a biological role are placed under a specific pathway. STRING-db was used to predict KEGG pathways and extract the predicted and experimental protein-protein interactions for the *C. acnes* KPA171202 proteome. KEGG orthology (KO) prediction resulted in the association of KO for 1503 CDS, and STRING-db associated KEGG pathways for 1593 proteins. In CAcnesDB, assignment of KEGG pathways was done for 687 additional proteins, summing up to KEGG pathway associations for 1596 proteins. Some examples are: (a) PPA0126 which encodes for ‘Putative penicillin-binding protein’ was assigned to three KEGG pathways terms: Peptidoglycan biosynthesis (pac00550), Metabolic pathways (pac01100), beta-Lactam resistance (pac01501), (b) PPA0130 which encodes for ‘Glycosyl transferase’ was assigned to Exopolysaccharide biosynthesis (pac00543) and Metabolic pathways (pac01100), (c) PPA1349 which encodes for ‘glutamate N-acetyltransferase (argJ)’ was assigned to five pathway terms: ‘Arginine biosynthesis (pac00240)’, ‘Metabolic pathways (pac01110)’, ‘Biosynthesis of secondary metabolites (pac01110)’, ‘2-Oxocarboxylic acid metabolism (pac01210)’, and ‘Biosynthesis of amino acids (pac01230)’.

Using the Predicted Prokaryotic Regulatory Proteins (P2RP) Webserver, the predicted regulatory proteins resulted in the identification of 12 histidine kinases and 12 response regulators. DeepTFactor employs three parallel convolutional neural networks and predicts if the sequence of a queried protein is a transcription factor. Using the DeepTFactor, 106 proteins were associated with a regulatory role. Using the VirulentPred server, the prediction of virulence resulted in 1,622 proteins that have a virulence association, as has been listed in Sheet S1 (Summary stats). A BlastP search (e-value ≤ 0.001) against set A and set B in the Virulent Factor Database (VFDB) resulted in 438 and 550 proteins having a virulence association.

### Enriching Genome Annotation with 3D structures

Colabfold was used to predict structures for the proteins coded by the *C. acnes* genome, which resulted in models for 2280 proteins. Supplementary Sheet S1 summarizes the results at each step of the structure-based annotation pipeline. The top-ranked predicted structures for each of the 2280 proteins constituted the *C. acnes*-structural proteome and were used for the prediction of the pocketome and associated ligands. The quality of the structural predictions from the ColabFold of the *C. acnes* proteome was analyzed. AlphaFold2, along with the actual three-dimensional coordinates, gives a measure of how confident one can be about the prediction itself, which is the predicted local distance difference test (pLDDT). Based on the reported AlphaFold thresholds, at the residue level, ColabFold generated high-confidence predictions for 67.4% of the *C. acnes* proteome, intermediate-confidence predictions for 18.9%, low-confidence predictions for 5.9%, and no prediction (i.e., pLDDT < 50) for 7.8% (Figure S4). To assign fold information for the modelled proteins (*n*=2280), a structural alignment was performed, where each of the 2280 proteins were aligned against non-redundant protein structures present in PDB (*m*=26,775). The program TM-align was used to establish the scan. This resulted in a 6,10,47,000 combination of *mn* pairs (2280 × 26775), which is roughly equivalent to 61 million pairs of structural alignments (see Supplementary Sheet S1,Summary stats). The top folds in the modeled *C. acnes* proteome include TIM beta/alpha-barrel, Rossmann(2×3)oid, Ferredoxin-like, OB-fold, UBA-type 3-helical bundle, etc. The top superfamilies in the modeled proteome include ‘Aldolase’, ‘Dihydropteroate synthetase-like’, ‘Phosphoenolpyruvate/pyruvate domain’ (Figure S5).

### *C. acnes* Pocketome

To predict possible small-molecule ligand binding pockets, three binding pocket prediction methods were used: Depth-based (i.e., PocketDepth), Energy-based (i.e., FPocket), and Voronoi tessellation-based (i.e., Site Hound). All three methods were run independently on each of the 2280 modelled structures (3 × 2280 = 6840). The numbers of pockets predicted by each of the methods are PocketDepth: 51,949, SiteHound: 22,800, and FPocket: 25,487. From these, consensus pockets (CP) were identified as described earlier (Figure S2). The CPs from the FPocket output are used for this analysis, and only those FPocket pockets with at least 4 residues that match the pockets predicted by PocketDepth and SiteHound were considered further. Based on the pLDDT scores, the percentage of high and intermediate confidence residues (pLDDT ≥ 70) with respect to the total number of residues in the pocket was calculated to evaluate the quality of the individual pockets. Among all the AlphaFold pockets, about 77.58% are located in protein structure regions with relatively high and intermediate confidence (ratio > 0.7). This resulted in 11,266 CPs spanning over 2034 proteins, which together constitute the *C. acnes*’ Pocketome.

### *C. acnes* predicted Metabolome, CApM

The Consensus Pockets (CPs) thus obtained are then compared to the known binding sites derived from the Protein Data Bank (PDB). Based on the analysis, the obtained CPs were recognized to have the binding capacity for the corresponding ligands associated with the structurally aligned PDB templates. Around 11,266 pockets exhibited significant similarity in the entire consensus pocketome to the known binding site in PDB, leading to the derivation of ligand associations for 1865 proteins (Figure 2A). The set of ligands associated with the pocketome is hereafter referred to as the predicted metabolome of *C. acnes* (CApM), which consisted of 1954 unique ligands associated with 1865 proteins. As the analysis has been conducted at the genome scale, the predicted ligands qualitatively comprise the predicted metabolome in *C. acnes* (CApM). Analysis of the CApM indicates that it captures most of the reported biochemical reactions in the literature. The coverage of the structural models and their associated ligands is represented in terms of KEGG pathways in Figure 2B. The *C. acnes* KEGG metabolic map shows the highlighted proteins that have modeled structures (in black edges) and ligands that have been associated with these proteins in Figure 2C. It draws out the extensive coverage of the identified ligands spanning over the cellular metabolism in *C. acnes*. Figure 2D shows the distribution of ligand hits obtained for the predicted sites in the pocketome. The ligands are ordered according to their molecular weight on the x-axis. In Figure 2E, the most frequently observed ligands are FAD, followed by NAD, ADP, HEM, NAP, and ATP. The predicted ligands that can bind to the *C. acnes* pocketome span a wide range of molecular weights, the lightest detected being acetate (ACY, 60.052) to one of the heaviest being Co-methylcobalamin (COB, 1344.382). Binding energies (ΔG) for all ligand–protein pairs were calculated using AutoDock. Among the 1865 proteins in CApM, 1562 exhibited binding energies lower than –5 kcal/mol with their respective ligands, suggesting a higher likelihood of stable ligand binding to these proteins.

**Figure 2:**
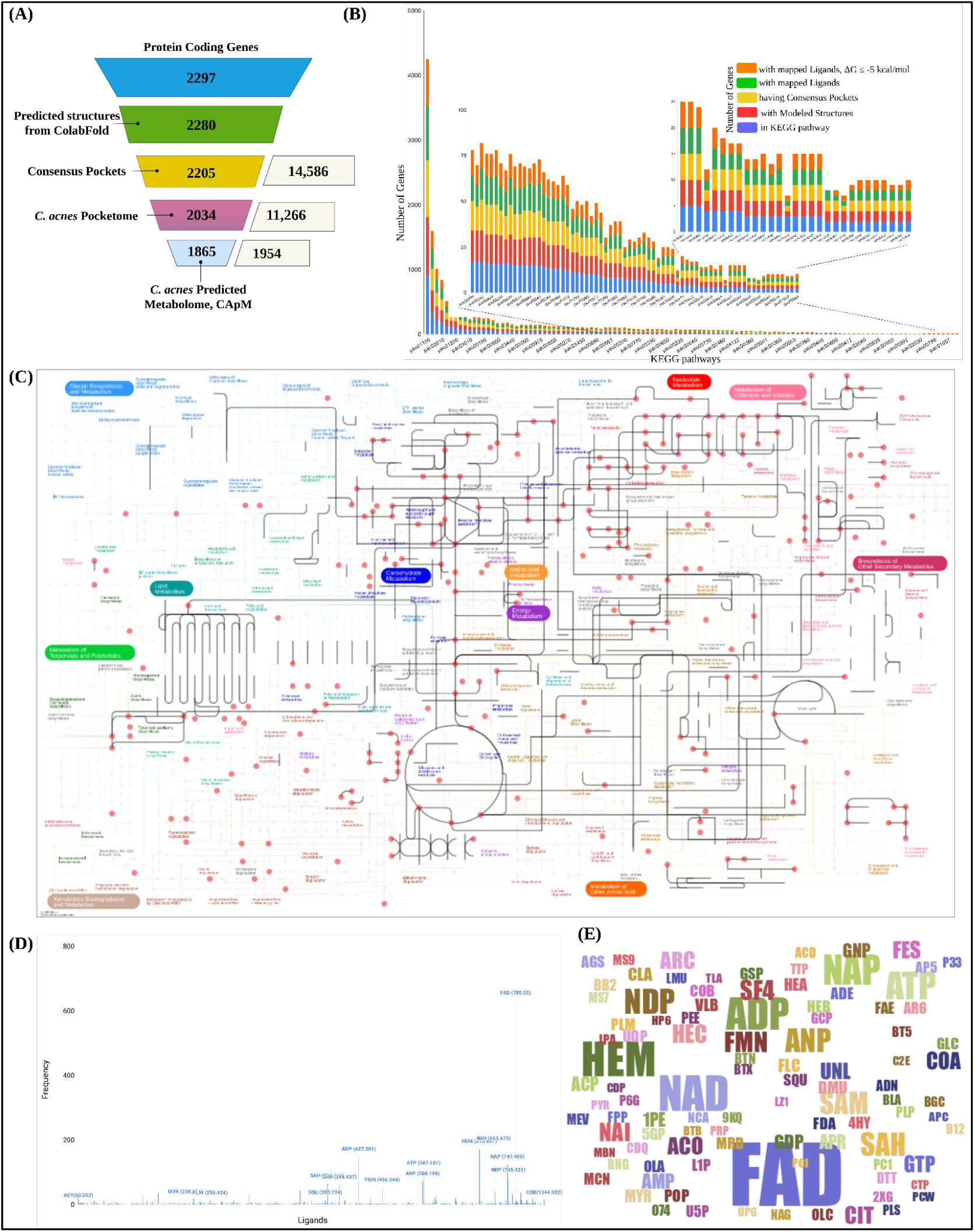
(A). The computational pipeline used for Proteome, Pocketome, and ligandome identification for the *C. acnes* interactome profile. The pipeline involved stepwise filters based on the modeled proteome, Consensus Pockets (CP) identification, and CApM analysis. (B) A stacked bar plot showing the coverage of protein structures and the confident ligand associations available with respect to the KEGG pathways. For each pathway, the lowermost bar in the stack corresponds to the number of genes or proteins in the pathway, the middle bar indicates the number of structural models available for the pathway, followed by the bars indicating the number of proteins for which ligand annotations are made based on the consensus pocketome and CApM, and the top-most bar indicates the number of proteins for which the mapped ligands have corresponding ΔG ≤ −5 kcal/mol. Each stack corresponds to one KEGG pathway in *C. acnes*. (C) Metabolic map of central metabolism in *C. acnes*, indicating extensive coverage of ligand annotation in the *C. acnes* predicted metabolome. The edges colored in black indicate the availability of protein structure catalyzing the reaction, and the nodes colored in red represent the small ligand molecules taking part in the reaction for which the binding site has been mapped onto the respective protein structure. The figure is generated using ipath3^35^. (D) An illustration of ligand associations for *C. acnes* Confidence Pocketome. The ligands are ordered by their molecular weights. The frequency on the Y-axis indicates the number of occurrences of the binding site of that ligand in the *C. acnes* pocketome. (E) A word cloud representing the top 100 most frequently occurring ligands from CApM of *C. acnes*. The size of the letters is directly proportional to the frequency of occurrence.

### Annotation-Quality

Annotations present the challenge of describing the annotation quality. This was dealt with by designing a scoring method to guide the annotation by identifying pieces of information and whether the assigned annotation is valid. Keeping this in mind, an annotation score was developed, where a value of ‘1’ was given if a particular piece of evidence was found to be making a connection with the given functional description; a score of ‘0’ was given for the lack of any such evidence, as outlined in Figure 3A. An example of computation of annotation-score for the assigned annotation in PPA0075 is shown in Figure 3B.

**Figure 3:**
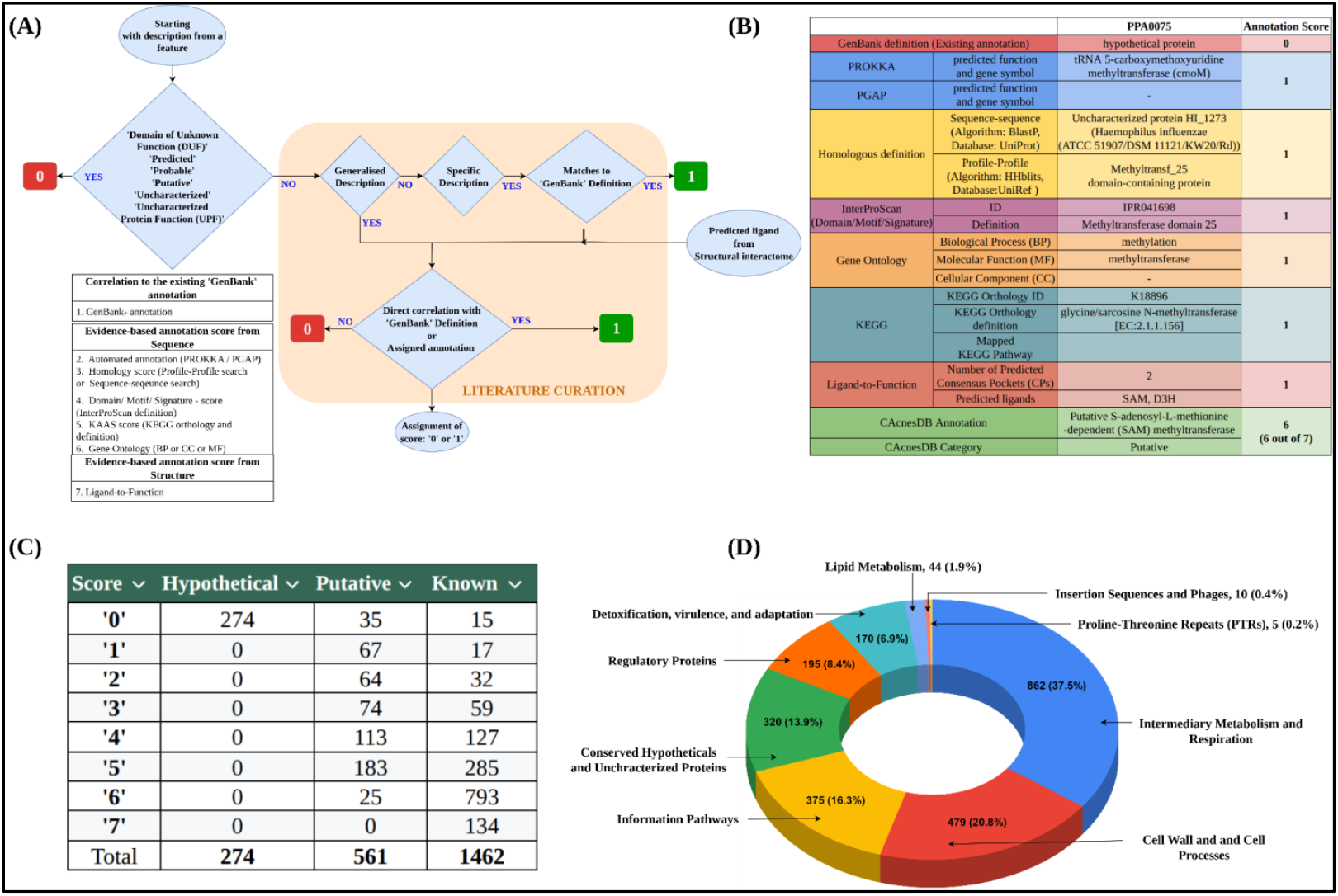
(A) Evidence-based annotation scoring schema. The figure illustrates the assignment of the annotation score to the revised/assigned annotation based on seven sections of the annotation pipeline. The steps were followed for each of the individual proteins in the proteome. Each step shows the assignment of either score ‘1’ or ‘0’ based on the match (presence/absence) to the particular section. Assessment of the annotation score was conducted with literature-based verification, wherever necessary. (B) Confidence-score assignment shown with an example of PPA0075; where it was annotated to be a ‘Putative SAM-dependent methyltransferase’ and was given a score of 6. (C) Distribution of annotation score across the three categories: Hypothetical, Putative, and Known. (D) Functional classification of *C. acnes* KPA171202 genes.

Summarization of the annotation score across the three categories showed 134 genes from the ‘Known’ category, which received an annotation score of ‘7’ (Figure 3C). This shows evidences from all seven categories (‘correlation with the existing annotation’, ‘homology searches’, ‘KEGG orthology’, ‘Gene Ontology’, ‘Domain-based correlation’, and ‘Ligand to function’) correlated with the assigned functional descriptions. The genes from the ‘Hypothetical’ category received a ‘Zero’ annotation score, showing the absence or no correlation of any evidence with these genes. The largest fraction, 789 genes from the ‘known’ category, received an annotation score of ‘6’, which underlines the rigorous steps followed in our annotation pipeline.

### Consolidated Annotations

The database, constructed specifically for *C. acnes* research, presents the consolidated annotation available for each protein. The protein-encoding genes were classified into nine functional classes (Figure 3D). These are (1) Intermediary metabolism and respiration (862/2297, 37.5%), (1) Cell wall and cell processes (479/2297, 20.8%), (3) Information pathways (375/2297, 16.3%), (4) Conserved hypotheticals and uncharacterized proteins (320/2297, 13.9%), (5) Regulatory protein (195/2294, 8.4%), (6) Detoxification, virulence, and adaptation (170/2297, 7.4%), (7) Lipid metabolism (44/2297, 1.9%), (8) insertion sequences and phages (10/2297, 0.4%), and (9) proline-threonine repeats (PTRs; 5/2297, 0.2%). The largest functional category was observed to be ‘Intermediary Metabolism and Respiration’, containing 862 enzymes, 766 of which were known and 96 were newly added/improved from our annotation. Some examples where annotation is enriched from our work are (i) PPA0083 is identified as ‘Putative 1,3-beta-galactosyl-N-acetyl hexosamine phosphorylase’ (as compared to the label of ‘hypothetical protein’ in the previous annotation), (ii) PPA2025 is ‘Putative FMN reductase (NADH) RutF’ (as compared to ‘hypothetical protein’ in the previous annotation), (iii) PPA1688 is ‘Putative phosphoheptose isomerase’ (as compared to hypothetical protein in the previous annotation). The different aspects of annotation for all the given examples are listed in Supplementary Sheet S2.

The ‘Cell wall and Cell Processes’, was another major functional category, that has all the ABC transporters, membrane permeases, and proteins facilitating cell divisions. This class includes 410 known and 69 new/improved annotations. Examples include (i) PPA0637 encoding ‘Putative DivIVA domain-containing protein’ as compared to ‘hypothetical protein’ in the previous annotation, (ii) PPA0231 is ‘Putative pilus assembly protein TadB’, as compared to hypothetical protein in the previous annotation). Another key functional category is ‘Information Pathways’, which includes genes essential for maintaining genetic information and supporting core cellular functions. This category encompasses genes involved in DNA modification and repair, as well as RNA subunits that play roles in replication, transcription, and translation.

Regulatory proteins, such as transcription factors, histidine kinases, and response regulators, play a central role in controlling the signaling pathways that enable the bacterium to adapt to environmental stress and regulate the expression of virulence factors. Genes involved in these adaptive responses, including those related to detoxification and virulence, are categorized under the functional class of ‘Adaptation, Detoxification, and Virulence’. Some examples are (i) PPA0389 is ‘Putative Asp23/Gls24 family envelope stress response protein’, as compared to ‘hypothetical protein’ in the previous annotation. The Asp23 family known to be present in acid-producing bacteria to maintain homeostasis in response to sudden pH changes, (ii) PPA1308 is now annotated as ‘Putative Universal stress protein A, UspA’ belonging to the ‘Universal stress protein family’ in Pfam with a ‘UspA domain’, and ‘Rossmann-like alpha/beta/alpha sandwich fold’ from domain analysis. (iii) PPA0653, previously labelled as ‘hypothetical protein’, now identified to be a Putative type II toxin-antitoxin (TA) system HicA family toxin’ of the HicA superfamily. (iv) PPA2046 is ‘Putative Trehalose utilization protein ThuA’ (as compared to hypothetical protein in the previous annotation).

The smallest portion of the genome, i.e., the PTR class, has five genes, which have been shown to be involved in colonization/adhesion/inflammation. PPA2127 has PT repeats at its C-terminus and was found to be Dermatan sulfate-binding adhesin 1 (DsA1), which is a host cell-surface attachment protein having dermatan-sulphate-binding activity and immunoreactive properties. PPA2210 and PPA2270 encode Dermatan sulfate adhesin (DsA2) and L,D-transpeptidase LppS-a putative secreted lipoprotein, respectively. PPA1715 was identified as TQXA, a thioester domain-containing protein. Dermatan sulfate/chondroitin sulfate (DS/CS) are skin glycosaminoglycans that mediate the adherence of bacteria to epithelial cells, including staphylococci, and participate in infections and mediate inflammatory response. Transposases (PPA0858, PPA0876, PPA1974, PPA2354, PPA2355, PPA2377, PPA2391) and conserved phage-associated proteins (PPA1069, PPA1594, PPA1604) are placed under the class for ‘insertion sequences and phages’.

Put together, our work has resulted in enriched annotations for 2072 proteins, of which 225 proteins were previously identified as hypothetical proteins, are assigned new functional annotations. Some of these newly annotated proteins belong to ‘Regulatory proteins’, examples being (a) PPA2204 encoding for ‘Putative transcriptional regulatory, WhiB’, (b) PPA0329 encoding for ‘Putative transcriptional regulator (RNA polymerase sigma factor) ‘. For example, (a) PPA1567, previously annotated as a hypothetical protein, our enhanced annotation identifies its closest homolog to be a membrane protein and Disulfide oxidoreductase D, the presence of a thioredoxin domain, associated with being an integral component of the membrane, and its overall function from the sequence analysis is established as ‘Putative thioredoxin’; (b) PPA1379, previously annotated as a hypothetical protein, our annotation identifies its closest homolog to be a sec-independent protein translocase protein tatA, presence of sec-independent protein translocase protein tatA domain, associated with biological process of transmembrane transport, having transporter activity; and hence is finally annotated as Putative sec-independent protein translocase protein TatA (Figure S1(b)).

Some examples of the new annotations from sequence- and structure-based cues are given in Supplementary Table S2. (i) PPA2105 is the ‘Triacylglycerol lipase (GehA)’ has 2 CPs, of which one is predicted to bind to Palmitic acid (PLM) with a binding energy of −7.5 kcal/mol (Figure 4D). (ii) Both PPA0075 and PPA0255 are annotated as ‘putative S-adenosyl-L-methionine-dependent (SAM) methyltransferase’, and have a CP each that can bind to S-Adenosylmethionine (SAM) with binding energies of −7.57 kcal/mol and −8.37 kcal/mol, respectively. As an example, the finding from the individual steps for the structure-to-function annotation in PPA0075 is depicted in Figure 4A. (iii) PPA2025 was characterized to be a ‘Putative FMN reductase (NADH) RutF’ and has 6 CPs, of which one has the capacity for binding Flavin Adenine Dinucleotide (FAD, ΔG= −3.32 kcal/mol). (iv) PPA1036, PPA1995, PPA2401 are characterized to be ‘Putative GNAT family N-acetyltransferases’, all having a CP predicted to bind to ACO (Acetyl Coenzyme *A (ACO) with calculated binding energies of −7.58 kcal/mol, −8.48 kcal/mol, and −11.93 kcal/mol, respectively. (v) PPA0410 is annotated to be ‘O-acetyl-ADP-ribose deacetylase’ which has 3 CPs, of which one is predicted to bind to ‘Adenosine-5-diphosphoribose’ (APR) with binding energy of −9.38 kcal/mol (Figure 4B). (vi) PPA1407 is annotated to be a ‘Putative type I Biotin synthase (BioB)’ and has three CPs, of which two have binding capacity for 4Fe-4S cluster (SF4) and S-Adenosyl-L-Homocysteine (SAH) with binding energies of −3.57 kcal/mol and −4.31 kcal/mol, respectively (Figure 4C). A recent study revealed that type II Biotin Synthase (BioB) in anaerobes binds 4Fe–5S clusters, unlike type I BioB in aerobes and facultative microbe genera, which binds 2Fe–2S and 4Fe-4S clusters^36^. (vii) PPA0731 was annotated to be a barstar-like family protein. Barstar is an intracellular repressor gene that binds and inhibits the extracellular ribonuclease barnase activity. Together, barnase and barstar mirror the toxin-antitoxin system (detected first in *Bacillus subtilis* strain H), which prevents the bacteria from digesting their own protein-synthesis machinery. (viii) PPA2334 is assigned as a PPi-type Phosphoenolpyruvate Carboxykinase (PPi-PEPCK), which catalyzes the formation of oxaloacetate from phosphoenolpyruvate and inorganic pyrophosphate. PPi-PEPCK is an alternative to GDP/ATP-based enzymes, and PPi-PEPCK has been shown to rewire the central-carbon metabolism in other Propionibacterium spp. (ix) PPA2204 is annotated as a ‘Putative transcriptional regulator, WhiB’ and has 2 CPs, having binding capacity for ATP and FAD. (x) Putative Phosphomannomutase from the HAD superfamily (PPA2023) is involved in the detoxification of phosphorylated compounds and pseudouridine. PPA2023 has 2 CPs with binding capacity for CIT (Citric acid), and PZH (1-(4-bromophenyl)methanamine. PPA0806 is annotated as a ‘Putative maleylpyruvate isomerase’, with 2 CPs, both predicted to bind FAD. (xi) PPA0047 is annotated as a LysM peptidoglycan-binding domain-containing protein. LysM-containing fusion proteins, such as Glucosaminidase (AcmA in *L. lactis*), endopeptidase (LytE), and muropeptidase (murO), by their binding and lytic activities, modulate host-microbe interactions. In addition to various lytic functions, lysM-containing proteins have antigenic and virulence properties. They are believed to be involved in bacterial adaptation to changing pH. (xii) PPA0231 is annotated as a ‘Putative pilus assembly protein TadB’, with 4 CPs, predicted to bind actinonin (BB2, a naturally occurring antibacterial agent; with a binding energy of −6.29 kcal/mol), Cholesterol (CLR, −8 kcal/mol), (3S,7S,11S)-3,7,11,15-tetramethylhexadecan-1-ol (ARC, ΔG = −5.17 kcal/mol), and heme (HEM, ΔG = −5.09 kcal/mol). (xiii) PPA0064 is annotated as a HAD hydrolase (family IA, variant 3) with 1 CP, predicted to bind 2-phosphoglycolic acid (PGA, ΔG = −1.84 kcal/mol). (xiv) PPA0883 is a ‘2-keto-3-deoxygluconate kinase’ with 3 CPs, of which one is predicted to bind to 2-keto-3-deoxygluconate kinase (KDG, ΔG = −3.63 kcal/mol). (xv) PPA0768 is a ‘Pseudouridine synthase’ which has 4 CPs, of which one is predicted to bind to Urindine-5’-monophosphate (U, ΔG = −8.48 kcal/mol). These examples illustrate the function association of the mapped ligands with their corresponding proteins. Details of their annotation is provided in the Supplementary Table S2.

**Figure 4:**
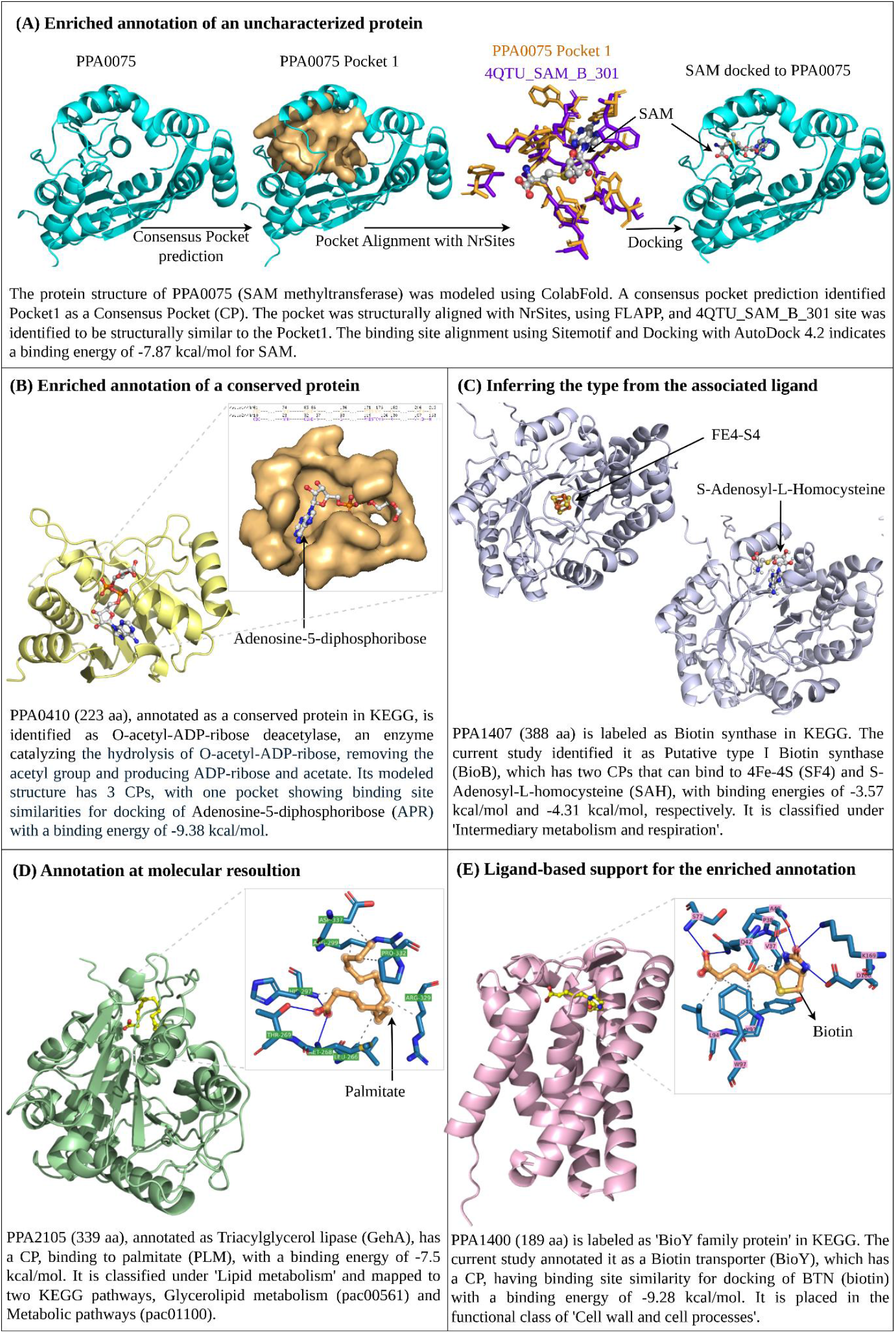
Different aspects of annotations are shown in the table, (A) from assigning the putative function to an uncharacterized protein (PPA0075), (B) to assigning a annotation for a conserved proteins (PPA0410), (C) inferring the type of enzyme from the associated ligand (PPA1407), (D) finding the substrate for the enzyme catalysis (PPA2105) and (E) confirming the existing annotation with the associated ligand (PPA1400).

### Transporters in *C. acnes*

*C. acnes* proliferates alongside the sebaceous glands present inside the pilosebaceous unit (PSU). It has saprophytic properties and acquires nutrients from the secretions of sebaceous glands, such as sebum lipids. The cell membrane is the interface between the intracellular metabolism and the external resources for nutrient availability. The bacterial cell membrane facilitates the host-pathogen interactions by mediating the exchange of metabolites, acquisition of nutrients, and excretion of wastes in its microenvironment. Bacterial ABC transporters, permeases, and protein exporters are the known mediators and facilitate metabolite exchange, contributing to host-pathogen interactions. This study has resulted in improved annotations for membrane transporters and their associated ligands that capture the probable array of small-molecule metabolites being exchanged between the host skin and the *C. acnes*.

From our annotation, out of the 359 transporters, 319 have mapped 521 unique ligands. The *C. acnes*’ transporters and the associated predicted metabolome are listed in the Supplementary Sheet S3. In all, these 359 transporters include 181 ABC transporters, 17 proteins involved in protein export, 236 proteins having transmembrane transport activity, and 188 proteins associated with transport activity. Many naturally occurring skin metabolites, sugars, and lipids, such as squalene, lipid fragments, glycerol, testosterone, and retinol, are found to be present in the transporters’ predicted metabolome. The following examples illustrate the newly added annotations: (a) PPA1400 is a Biotin transporter, having a CP that can bind to Biotin with a binding energy of −9.28 kcal/mol (Figure 4E). (b) PPA1936 is an ABC transporter ATP-binding protein that can bind to arachidonic acid with a binding energy of −7.2 kcal/mol. (c) PPA0016 is an ABC transporter permease that has a pocket predicted to bind SQU (2,10,23-trimethyl-tetracosane)-a lipid fragment with a binding energy of −8.1 kcal/mol. (d) PPA1966 is the Glucose/mannose transport system substrate-binding protein, which has CPs having binding-site similarities for carrying GLC (alpha-D-glucopyranose), HHW ((∼{E})-3-[3-[(4-chlorophenyl)carbamoyl]phenyl]prop-2-enoic acid), and FAD, with binding energies of − 5.17 kcal/mol, −6.09 kcal/mol, and −5.68 kcal/mol, respectively. (e) PPA0657 is a Heme ABC superfamily ATP-binding cassette transporter that has a pocket predicted to bind HEM (Protoporphyrin IX containing Fe), SF4 (4Fe-4S cluster), with binding energies of −7.29 kcal/mol and −3.2 kcal/mol, respectively. Additionally, it has other pockets that can possibly bind a lipid fragment (SQU) and sulfoquinovosyldiacyl-glycerol (SQD) with binding energies of −7 kcal/mol and −6.11 kcal/mol, respectively.

### CAcnesDB Database Functionalities

The CAcnesDB home page provides access to various aspects of annotation, namely (i) Protein Details, (ii) Domain and Gene Ontology, (iii) Protein Structure and Binding Pockets, (iv) Protein-Protein Interaction (v) Pathway Information. The user can search for the listed proteins through the listed gene symbols, UniProt IDs, NCBI IDs, or assigned annotations. CAcnesDB presents modeled protein structures (Colabfold predicted top-scoring modeled structure for each query protein) for 2,280 protein sequences that can be assessed in the ‘Protein Structure and Binding Pockets’ tab. In addition, the tab provides the list of predicted CPs for the queried protein (if any). The user can select and visualize the pockets in the interactive graphics interface (Figure S6). An interface has also been provided, integrating the available data from other external resources such as STRING and KEGG.

## Discussion

Genome annotation is an iterative activity and periodic updation is necessary to benefit from the knowledge that is continuously being accumulated in databases and primary literature for many organisms. A number of draft genome sequences of *C. acnes* strains have been reported, indicating the interest in studying this organism, and we expect our annotation reported here to be widely useful.

The annotation pipeline devised by us uses a combination of automated and manually curated updates of the whole genome characterization. The enriched annotations involve various aspects of gene characterization, which were extracted from both the sequence and structure-based analyses. While the sequence-based methods provided functional descriptions in terms of Gene Ontology (BP, MF and CC) classes, KEGG pathways, virulence and regulatory role predictions, the structure-based methods provided a deeper layer of evidence, by identifying metabolites -possibly associated with each pocket, defined with the exact amino acid residues lining them, at a proteome scale.

Proteins function in dynamic networks to regulate processes like signaling, metabolism, and cell adhesion. Protein–protein interaction information aids in functional annotation and pathway mapping, metabolic enzymes support genome-scale metabolic model reconstruction, and regulatory proteins help elucidate regulatory networks. Previously identified lipases, endoglycoceramidases, sialidases, neuraminidases, and pore-forming factors, genes involved in anaerobic respiration and adaptation aided in understanding the pathogenic capabilities of *C. acnes*, and our work enriches the annotation of these, adding consolidated information on likely amino acids at the pockets, ligands, protein-protein interactions and functional categories. In addition to the annotation of many previously uncharacterized proteins, it provides new annotations to a few hundreds and enriches the annotation of a large number of proteins. For example, Triacylglycerol lipase (GehA), classified under Lipid metabolism, was mapped to the KEGG pathways for Glycerolipid metabolism (pac00561) and general Metabolic pathways (pac01100) and has been identified to have binding sites for palmitate (PLM). Obligate anaerobes have type II Biotin synthases (BioB), a radical S-adenosyl-L-methionine enzyme, which are known to bind to an auxiliary 4Fe–5S cluster, instead of 2FE-2S and 4FE-4S clusters in aerobes and facultative bacterial genera^36,37^. The predicted binding of a 4FE-4S cluster (SF4) and S-Adenosylmethionine (SAM) in *C. acnes* BioB suggests a type I biotin synthase, and warrants further experimental and biochemical validation. BioB is classified under intermediary metabolism and respiration, and is mapped to three KEGG pathways: biotin metabolism (pac00780), which includes two KEGG modules (pac_M00123 and pac_M00577), general metabolic pathways (pac01100), and biosynthesis of cofactors (pac01240). Many of the ABC transporters (pac02010) and membrane permeases that were placed under the functional class of ‘cell wall and cell processes’ were found to have pockets that can bind to various skin-related metabolites, such as squalene (SQL), sebum lipids like oleic acid (OLA), myristic acid (MYR), and vitamins. The knowledge of the specific substrates that can bind to the transmembrane transporters and permeases serves as hypotheses to study metabolite-centric host-pathogen interactions between *C. acnes* and human skin.

One of the characteristic features of *C. acnes* is having PT repeats (PTRs) that possibly have immunogenic properties, similar to the PPE repeats in *Mycobacterium tuberculosis*. Previous reports (McLaughlin et al.,^4^) have indicated PPA2210, PPA1715, PPA1879, PPA1880, PPA1881, PPA1906, PPA2130, and PPA2270 to contain PTRs. From our annotation, we have grouped five proteins in the PTR class, namely, PPA2127, PPA2210, PPA1880, PPA1715, and PPA2270. These genes encode adhesins (DsA1, DsA2), a secreted lipoprotein, and have also been placed under the class of ‘Adaptation, Detoxification, and Virulence’, due to their involvement in inflammatory responses and infections. Given that *C. acnes* phages exhibit a narrow host range, including strains such as *C. acnes* KPA171202^38^, conserved phage-associated proteins and transposases were classified under the category of ‘insertion sequences and phages’. Future studies may uncover additional genes belonging to this functional class, allowing a greater coverage of phage-related elements in the *C. acnes* genome. The multi-modal approach employed in the current study, including pathway mapping, domain and fold identification, and metabolite prediction, provides a robust foundation for decoding the remaining 13.9% of uncharacterized proteins and for uncovering additional functions within the annotated proteome. These resources are crucial for advancing systems-level understanding of *C. acnes* biology.

In conclusion, the genome annotation of *C. acnes* has been revisited using a multi-analysis pipeline, resulting in annotation for 2297 proteins with significant improvement in 2072 proteins. The analyses and the annotations are presented in CAcnesDB (https://proline.biochem.iisc.ac.in/CAcnesDB/), an online database providing integrated access to genome sequence, structural proteome, and literature curation for *C. acnes*. CAcnesDB stores hierarchical functional and structural characterization, i.e., modeled proteome, protein-protein interaction network, pocketome, and metabolome. To enable wide use of these data, CAcnesDB provides a suite of tools for searching, browsing, analyzing, and downloading the data. CAcnesDB also serves as a platform to add future genome annotations of related strains and perform a comprehensive analysis. By integrating a wide range of genomic data with tools for their use, CAcnesDB is a unique platform for both basic science research in *C. acnes*-related skin infections and disorders, as well as research into the discovery and development of drugs and possibly vaccines for acne vulgaris.

## Supporting information

Supplementary Sheet S1

Supplementary Sheet S2

Supplementary Sheet S3

Supplementary Method Doc1

Supplementary Results Doc2

## SUPPLEMENTARY

1. A list of tools and databases used for the functional annotation based on sequences, from structures, and the construction of CAcnesDB is provided in Supplementary Sheet S1 (Tools and Databases). For each step, summary statistics are written in the adjacent sheet in Supplementary Sheet S1 (summary stats).
2. A detailed sequence-to-annotation (homology prediction, domains/motifs/signatures prediction, gene ontology prediction, KEGG pathways association, virulence and regulatory role analysis), structure-to-annotation (SCOP fold, superfamily, and family assignment), assessment of the annotated function description (manual curation, literature-based up-to-date annotation, computation of annotation score) are provided in Supplementary Methods Doc1.
3. The distribution of GO terms (BP, CC, MF), quality of modeled proteome, and the SCOP-FOLD, family and superfamily across the *C. acnes* KPA171202, is provided in Supplementary Results Doc2.
4. The different aspects of annotation for all the given examples are reported in Supplementary Sheet S2.
5. The whole genome annotation for *C. acnes* KPA171202 carried out in CAcnesDB is provided in Supplementary Sheet S3. The transporters in *C. acnes* (ABC transporters, transmembrane permeases and the protein export) along with their mapped ligands, are listed in Supplementary Sheet S3.

## Author contributions

Conceptualization: B.P. and N.C.; Database consolidation: B.P., D.R., and N.R.; Methodology: B.P., R.S., and N.C.; Pipeline development: B.P., R.S., and N.C.; Processed computational analysis: B.P., S.S. and S.V.; Figure preparation: B.P., N.R.; Writing—original draft preparation: B.P., and N.C.; Writing—review and editing: B.P. and N.C.

B.P., R.S., and N.C. contributed to the discussion of all contents. B.P. coordinated the study, R.S. and N.C. advised research, and revised the manuscript. All authors participated in writing this manuscript.

## Acknowledgements

This work has been supported in part by grants from the Department of Biotechnology, Bioinformatics Centre (BT/PR40186/BTIS/137/3/2020) and (BT/PR40316/BTIS/137/83/2023). B.P. is grateful to the Ministry of Education for the research fellowship. Saurabh Dey is acknowledged for the help with manual curation and exploring formats for database population. N.R. is grateful to the Centre for Networked Intelligence (a Cisco CSR initiative) of the Indian Institute of Science, Bengaluru, for the research fellowship.

