## Supplementary Method Doc1 for "CAcnesDB: A database of *Cutibacterium acnes* with Integrated Functional Insights derived from Multi-modal Genome Annotation"

### Supplementary Methods

#### Sequence-to-Function Association

##### 1. Homology Prediction

Universal Protein Resource (UniProt) and its knowledge base (UniProtKB) are highly used EMBL-EBI repositories[1]. The UniProtKB contains (i) unreviewed protein sequences having computationally generated, large-scale functional annotations (TrEMBL) and (ii) reviewed protein sequences having high-quality, manually annotated, and non-redundant sequences (Swiss-Prot). UniProt Reference Clusters (UniRef) stores sets of protein sequences that share 50% (UniRef50), 90% (UniRef90), and 100% (UniRef100) identity[2]. Sequence-sequence homology search with the BlastP algorithm was performed for individual protein sequences by querying against the Swiss-Prot/Reviewed database. Profile-profile homology search employing HHblits (number of iterations = 2) was carried out for each protein against the UniRef100[3]. The annotation of the significant top hits (e-value < 0.001), resulting from both searches, was associated with the given query sequence.

##### 2. Domains/ Motifs/ Signatures Prediction

Protein sequences for individual genes were retrieved from KEGG (Kyoto Encyclopedia of Genes and Genomes) and subjected to a command-line version of InterProScan for domain-based functional characterization[4,5]. InterProScan gives a summarized functional annotation by combining individual matching signatures from each search application. It has 14 member databases, which work based on homologous/orthologous prediction.

##### 3. Gene Ontology Prediction

Protein sequences were subjected to Gene Ontology (GO) analysis employing Pannzer2. Pannzer2 is a protein homology search-based web-server tool for rapid functional annotation of proteins[6]. Along with GO terms, Pannzer2 also outputs functional descriptions for the queried protein sequences.

##### 4. KEGG Pathways Association

The Bio.KEGG.REST module was used to assign KEGG gene IDs to each gene based on matching gene coordinates. KEGG functionalities such as the KEGG Mapper and KEGG Reconstruct pathway tool were used for gene mapping and visualization in KEGG pathways[7]. The STRING database was used to add the generated KEGG pathways for the *C. acnes* KPA171202 proteome.

The systematic annotation pipeline based on the protein sequences is illustrated with an example for the gene PPA1379 in Figure S1(b). (1) Gene prediction resulting from PROKKA (Complement 15011567...1501390) and PGAP (Complement 1501031...1501294) indicated that PPA1379 encodes for tatA. The existing GenBank definition from NCBI showed it to be a hypothetical protein. (2) Domain/Motifs identification using InterProScan showed that the sequence of PPA1739 has the mttA/Hcf106 family (PF02416) domain. (3) Gene Ontology Prediction using Pannzer2 and InterProScan showed that the Biological Process (BP), Molecular Function (MF), and Cellular Component (CC) are protein transport by Tat complex, protein transmembrane transporter activity, and Tat protein transport complex, respectively. (4) Sequence-sequence homology search and Profile-Profile homology search results showed the closest homolog is a gene encoding for sec-independent protein translocase protein tatA. (5) KEGG pathway association using KEGG Mapper showed that the gene can be assigned to pac03070: Bacterial secretion system and pac03060: Protein export. Thus, based on the annotation from the sequence, gene PPA1379 was characterized as a sec-independent protein translocase protein, tatA.

#### Virulence and Regulatory Role Analysis

Virulence prediction was performed using VirulentPred and Virulent Factor Database (VFDB)[8,9]. VirulentPred predicts virulent proteins using a two-layer cascaded support vector machine (SVM). The initial layer trains and enhances the features of protein sequences, such as dipeptide composition (DPC), position-specific scoring matrices (PSSMs), and amino acid composition (AAC). The refined features are subjected to the second SVM layer for the final prediction[8]. VFDB provides experimentally characterized (set A) and predicted virulent factors (set B) in prokaryotes[9]. A sequence similarity search was performed using the BlastP algorithm to identify virulent proteins based on the closest match (e-value < 0.001) in the VFDB database (in both set-A and set-B). The union of all the predicted proteins from the virulentPred and VFDB databases was considered virulent. For the regulatory role prediction, P2RP and DeepTFactor were used[10,11]. P2RP is a web-based framework designed to identify prokaryotic regulatory proteins. It predicts Histidine kinases, Response regulators, Phosphotransfer proteins, Transcriptional Regulators, Sigma Factors, and other DNA-binding proteins. DeepTFactors is a deep learning-based framework used to predict transcription factors.

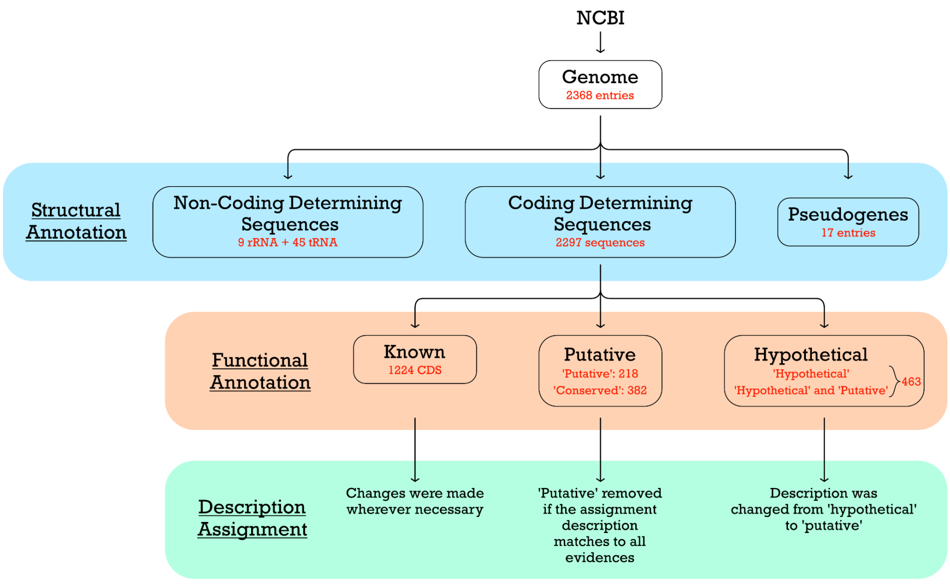
NCBI
Genome
2368 entries
Structural Annotation
Non-Coding Determining Sequences
9 rRNA + 45 tRNA
Coding Determining Sequences
2297 sequences
Pseudogenes
17 entries
Functional Annotation
Known
1224 CDS
Putative
'Putative': 218
'Conserved': 382
Hypothetical
'Hypothetical'
'Hypothetical' and 'Putative'
463
Description Assignment
Changes were made wherever necessary
'Putative' removed if the assignment description matches to all evidences
Description was changed from 'hypothetical' to 'putative'

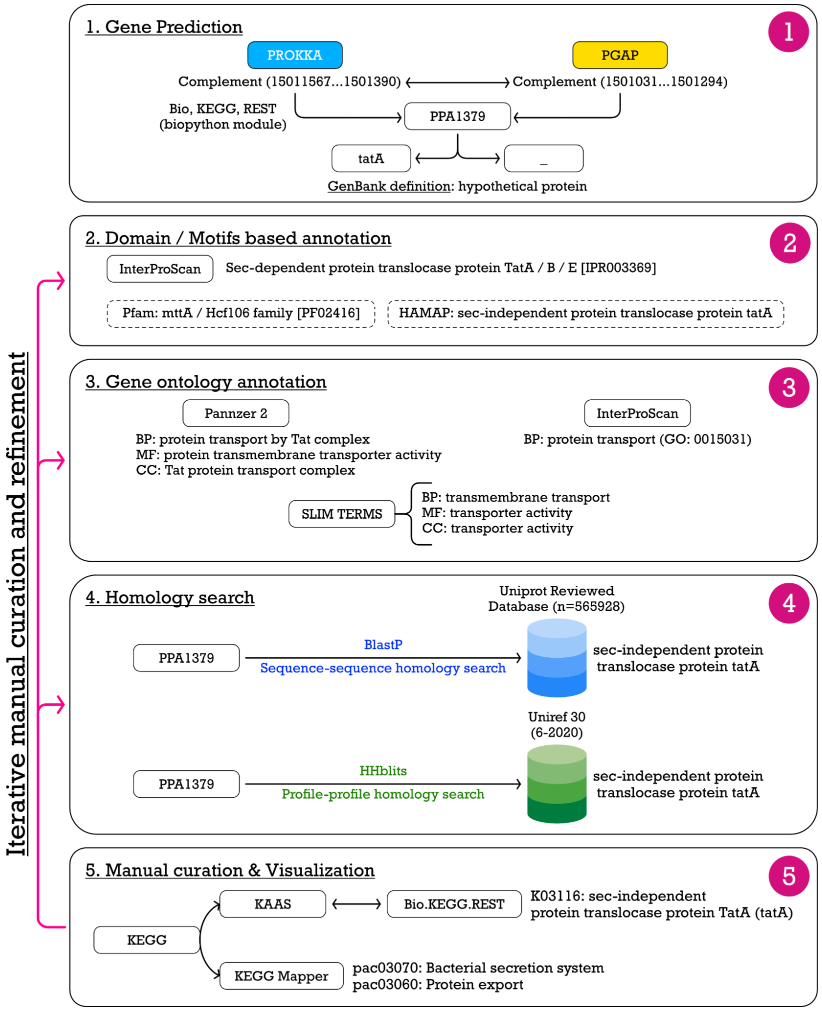
Iterative manual curation and refinement
1. Gene Prediction
1
PROKKA
PGAP
Complement (15011567...1501390)
Complement (1501031...1501294)
Bio, KEGG, REST (biopython module)
PPA1379
tatA
_
GenBank definition: hypothetical protein
2. Domain / Motifs based annotation
2
InterProScan
Sec-dependent protein translocase protein TatA / B / E [IPR003369]
Pfam: mttA / Hcf106 family [PF02416]
HAMAP: sec-independent protein translocase protein tatA
3. Gene ontology annotation
3
Pannzer 2
BP: protein transport by Tat complex
MF: protein transmembrane transporter activity
CC: Tat protein transport complex
InterProScan
BP: protein transport (GO: 0015031)
SLIM TERMS
BP: transmembrane transport
MF: transporter activity
CC: transporter activity
4. Homology search
4
Uniprot Reviewed Database (n=565928)
PPA1379
BlastP
Sequence-sequence homology search
sec-independent protein translocase protein tatA
Uniref 30 (6-2020)
PPA1379
HHblits
Profile-profile homology search
sec-independent protein translocase protein tatA
5. Manual curation & Visualization
5
KEGG
KAAS
Bio.KEGG.REST
K03116: sec-independent protein translocase protein TatA (tatA)
KEGG Mapper
pac03070: Bacterial secretion system
pac03060: Protein export

**Figure S1 (a):** Workflow for gene calling (structural annotation), initial segregation, and after the revised annotation, categorizing the genes into 'known', 'Putative', and 'Hypothetical' groups. (b) An annotation pipeline for annotation based on the sequence used for the individual proteins in the proteome. Each step shows different types of functional association, followed by iterative manual curation and assignment of the KEGG pathway to the associated function. The annotation pipeline is shown with an example for PPA1379.

### Other aspects of annotation

#### FOLD, Superfamily, and Family assignment.

Protein structures are highly informative and can be used to predict folds associated with a modeled structure. Known folds can provide supporting information for the existing annotation. SCOP2 is a database that stores the manually curated structural domains (Class, Fold, Superfamily, Family) for the structures available in the PDB database[12]. The 2280 proteins' predicted structures were aligned against non-redundant protein structures present in PDB (m=26,775). The TMalign program was used to establish scan[13]. It takes the predicted structures (atomic coordinate file) of individual proteins and outputs the aligned structures with alignment length, RMSD, sequence identity, TMScore1, and TMScore2. The scan resulted in 61047000 pairs of modeled structures with the PDB templates. Further, the results were filtered with different cutoffs, and the coverage of the modeled proteins was calculated. (i) TMScore1 and TMScore2 $\geq$ 0.7 [1023/2280 proteins] (ii) TMScore1 and TMScore2 $\geq$ 0.6 [1294/2280 proteins] (iii) TMScore1 and TMScore2 $\geq$ 0.5 [1546/2280 proteins]. 1546 PDB templates resulting from the TMScore1 and TMScore2 $\geq$ 0.5 were used to map to the available FOLD, superfamily, and Family in the SCOP2 database.

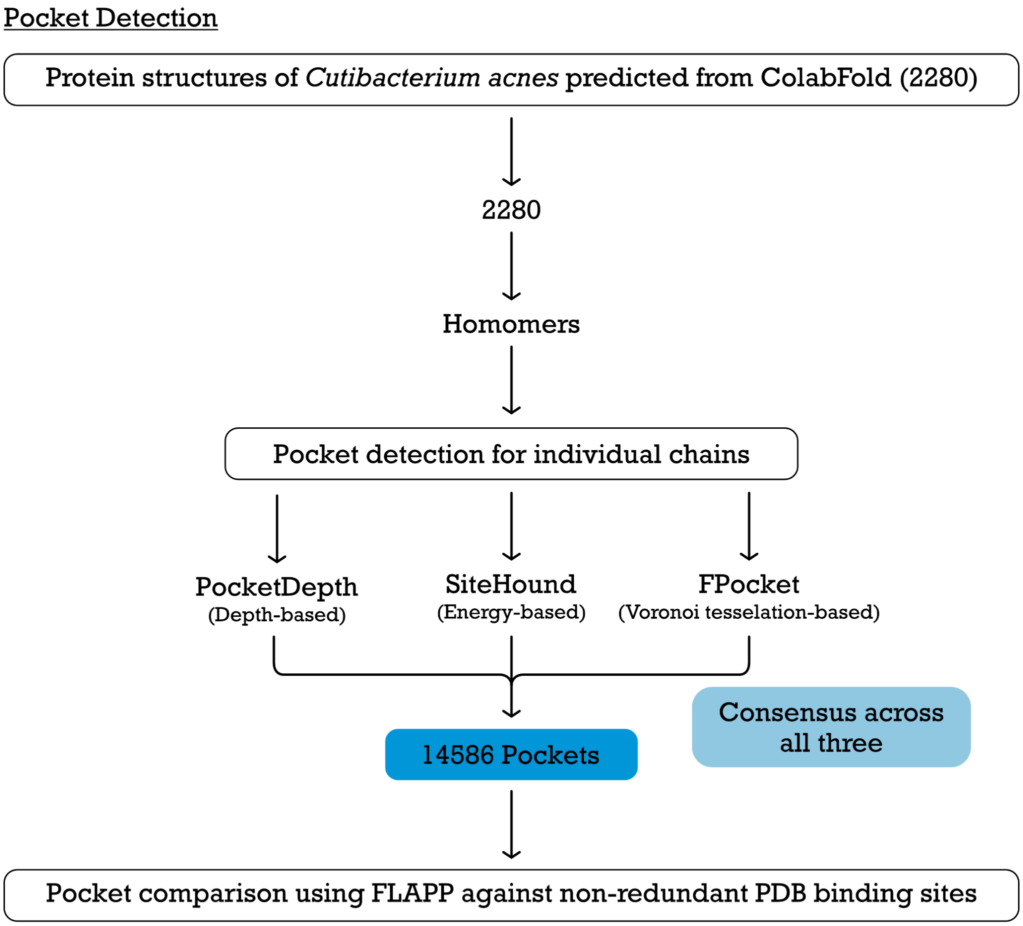


**Figure S2:** Workflow illustrating the functional annotation from structures (A) Modeling of the proteome in *C. acnes*; (B) Identification of ligand binding pockets from PDB. Three different algorithms were used for detecting pockets, and only those that were detected by all three were considered for further analysis. (C) Comparison of ligand binding pockets with the closest homolog (template) from PDB; Mapping the ligand from the template to the predicted model of each protein, and construction of ligandome in *C. acnes*. and correlating the associated protein function with the mapped ligand. PDB, Protein Data Bank.

### Assessment of the annotated functional description

#### 1. Manual Curation

Manual curation was iteratively carried out at each step of the pipeline. The results from individual steps were cross-checked with the 'GenBank' definition. In the case of indirect correlations, no functional description was assigned.

#### 2. Literature-based up-to-date annotation

Enormous progress has been made in the understanding of *C. acnes* biology since the sequencing and annotation of the reference strain KPA171202. These pieces of information from the literature are incorporated into the annotation pipeline.

#### 3. Computation of Annotation-score

A binary-score schema was developed to evaluate the confidence of the assigned annotation. The flowchart for the computation of the annotation score is depicted in Figure 3A.

i) The predicted function from each tool was manually checked, and a score '1' was given to the categories for which (a) direct correlation was present from a particular piece of evidence, (b) In case of multiple evidences, a Boolean logic of 'either' 'or' was followed. (c) If a particular description from an evidence suggests the possibility of correlation with a generalized term (d) Based on string match to the web search.

ii) Score '0' was given to the categories for (a) the absence of any evidence, (b) any indirect correlation between the evidence and the assigned annotation, (c) based on the string match to 'putative' or 'uncharacterized' or 'Protein with the domain of unknown function (DUF)'. (c) and in case of 'indecision' or 'doubt'.

iii) A special advantage was given to the literature-driven, experimentally characterized proteins. A score of '5' was assigned if the characterized protein has literature-based experimental validations. For example, gene PPA1939 encodes the unique antioxidant-radical oxygenase of *Propionibacterium acnes* (RoxP)[14], and the genes in the cluster PPA0859-PPA0865 are involved in Cutimycin biosynthesis[15].

An example of the computation of the annotation score for the assigned annotation is depicted in Figure 3B.

(4) Kanehisa, M. KEGG: Kyoto Encyclopedia of Genes and Genomes. *Nucleic Acids*

*Research* **2000**, *28* (1), 27–30. https://doi.org/10.1093/nar/28.1.27.
(5) *InterProScan 5: genome-scale protein function classification | Bioinformatics | Oxford Academic*. https://academic.oup.com/bioinformatics/article/30/9/1236/237988 (accessed 2025-07-23).
(6) *PANNZER2: a rapid functional annotation web server | Nucleic Acids Research | Oxford Academic*. https://academic.oup.com/nar/article/46/W1/W84/4993786?login=true (accessed 2025-07-23).
(7) Kanehisa, M.; Sato, Y.; Kawashima, M. KEGG Mapping Tools for Uncovering Hidden Features in Biological Data. *Protein Sci* **2022**, *31* (1), 47–53. https://doi.org/10.1002/pro.4172.
(8) Sharma, A.; Garg, A.; Ramana, J.; Gupta, D. VirulentPred 2.0: An Improved Method for Prediction of Virulent Proteins in Bacterial Pathogens. *Protein Sci* **2023**, *32* (12), e4808. https://doi.org/10.1002/pro.4808.
(9) *VFDB: a reference database for bacterial virulence factors | Nucleic Acids Research | Oxford Academic*. https://academic.oup.com/nar/article/33/suppl_1/D325/2505203 (accessed 2025-07-23).
(10) Barakat, M.; Ortet, P.; Whitworth, D. E. P2RP: A Web-Based Framework for the Identification and Analysis of Regulatory Proteins in Prokaryotic Genomes. *BMC Genomics* **2013**, *14*, 269. https://doi.org/10.1186/1471-2164-14-269.
(11) Kim, G. B.; Gao, Y.; Palsson, B. O.; Lee, S. Y. DeepTFactor: A Deep Learning-Based Tool for the Prediction of Transcription Factors. *Proceedings of the National Academy of Sciences* **2021**, *118* (2), e2021171118. https://doi.org/10.1073/pnas.2021171118.
(12) *SCOPe: improvements to the structural classification of proteins – extended database to facilitate variant interpretation and machine learning | Nucleic Acids Research | Oxford Academic*. https://academic.oup.com/nar/article/50/D1/D553/6447236?login=true (accessed 2025-07-23).
(13) Zhang, Y.; Skolnick, J. TM-Align: A Protein Structure Alignment Algorithm Based on the TM-Score. *Nucleic Acids Res* **2005**, *33* (7), 2302–2309. https://doi.org/10.1093/nar/gki524.
(14) *A novel enzyme with antioxidant capacity produced by the ubiquitous skin colonizer Propionibacterium acnes | Scientific Reports*. https://www.nature.com/articles/srep36412 (accessed 2025-07-23).
(15) *A Cutibacterium acnes antibiotic modulates human skin microbiota composition in hair follicles | Science Translational Medicine*. https://www.science.org/doi/10.1126/scitranslmed.aay5445?url_ver=Z39.88-2003&rfr_id=ori:rid:crossref.org&rfr_dat=cr_pub%20%200pubmed (accessed 2025-07-23).
