## Supplementary Results Doc2 for "CAcnesDB: A database of *Cutibacterium acnes* with Integrated Functional Insights derived from Multi-modal Genome Annotation"

#### Annotation Overview and Summary Statistics

| | KEGG (on 01/08/2025) | ∗UniProt (on 1/8/2025) | ∗NCBI (GenBank,RefSeq) | TBdb (on 1/8/2025) | †CAcnesDB (on 1/8/2025) |
|---|---|---|---|---|---|
| **Protein coding genes (CDS)** | 2297 | 2295 (299 reviewed + (1996 unreviewed) | 2425 | 2297 | 2297 |
| **Pseudogenes** | 17 | - | 88 | - | 17 |
| **Protein non-coding genes** | 54 (9 rRNA + 45 tRNA) | - | 76 (18 riboswitch, 1 RNAse_P_RNA, 9 rRNA, 1 SRP_RNA, 2 tmRNA, 45 tRNA | - | 55 (6 rRNA + 47 tRNA + 2 tmRNA) |
| **Protein with definite functional assignments (absence of ' Putative/ Probable/ Predicted/ Hypothetical')** | 1344 | 1824 | 1997 | 1208 | 1826 |
| **Protein with 'Probable/ Putative/ Predicted' functional assignments** | 472 (468 'putative'+ 16 'probable'+12 'predicted') -(24 'hypothetical') | 114 | 3 | 441 (444 'putative'+ 10 'probable'+11 'predicted')- 24 'hypothetical') | 215 (194 'putative' + 14 'probable'+ 7 'predicted') |
| **Hypothetical/ Uncharacterized/ Domain of Unknown (DUF) proteins** | 481 | 357 | 425 | 648 | 256 (225 are newly annotated) |
| **Protein with gene symbols** | - | 442 | - | 58 | 823 |
| **Protein with assigned domains** | 2085 (Pfam) | 1981 (Pfam, InterPro) | | 1841 (Pfam, Tigrfam) | 2085 (InterPro, Pfam, Gene3D, CDD, Coils, HAMAP, PRINTS, PrositePatterns, PrositeProfiles, SFLD, SMART, superfamily, Tigrfam, MobiDB, Panther) |
| **Protein with assigned Gene Ontology (GO) (BP or MF or CC)** | - | 1759 | 1267 | - | 1686 |
| **Protein with assigned KEGG orthology (KO)** | 1397 | - | - | - | 1503 |
| **Protein with assigned KEGG pathways/modules** | 909 (KEGG pathways + KEGG modules) | - | - | - | 1596 (KEGG pathways + KEGG modules) |
| **Structural models for proteins** | - | 2286 (AlphafoldDB) | - | - | 2280 (predicted models from ColabFold) |
| **SCOP domains** | - | - | - | - | 1205 (Superfamily and/or FOLD and/or Family) |
| **Binding ligands (predicted)** | - | - | - | - | 1865 |
| **Binding ligands with intermolecular binding energy <= -5 kcal/mol** | | - | - | - | 1562 |
| **Availability of Functional for the whole genome of *C. acnes*** | - | - | - | - | Yes |
| **Functional Classes** | - | - | - | - | 9 Functional Classes |
| **Availability of Annotation-confidence schema** | - | Yes | - | - | Yes |

† Manually curated, known non-redundant dataset

∗ Information retrieved in original form without redundancy filtering

Table S1: Summary statistics of the availability of results from each analysis carried out in the annotation of the *C. acnes* KPA171202. The results are compared with the available data from the other public resources, i.e., KEGG, UniProt, NCBI, and TBDB.

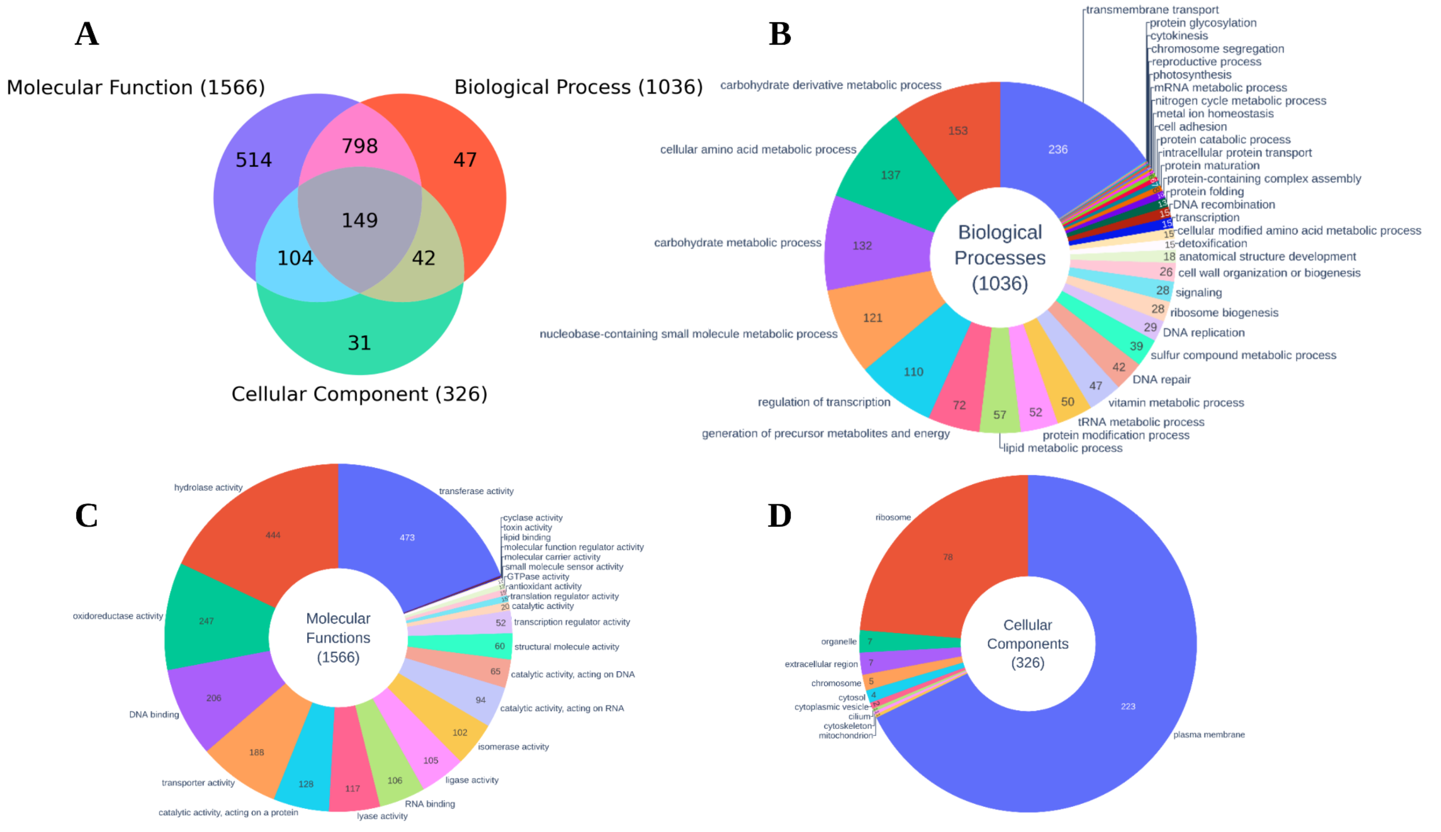


Figure S3: Pie chart representing the Gene Ontology (GO) analysis from Pannzer2 (a) Association of GO terms for *C. acnes* KPA171202 genome in three principal GO categories (BP: Biological Process, MF: Molecular Function, CC: Cellular Component). (b) The distribution of GO terms for the BP category across 1036 genes. (c)The distribution of GO terms in the MF category for 1566 genes. (d) The distribution of GO terms in the CC category for 326 genes.

### Structure-based Analysis

##### Modeled proteome's quality assessment

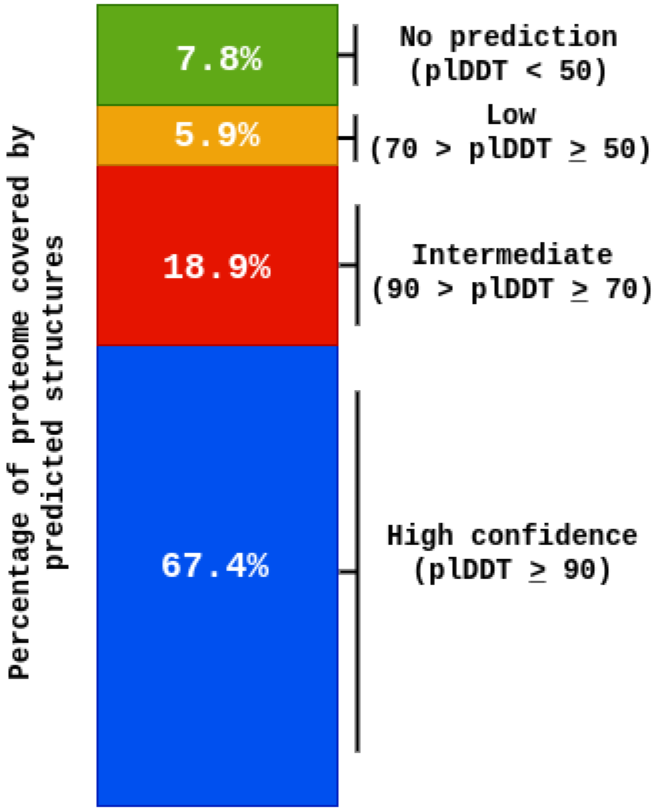


**Figure S4:** Coverage of the proteome by different ColabFold pLDDT score thresholds.

##### Summary of the associated FOLD, superfamily, and family

Unlike traditional homology modeling methods, AlphaFold does not give other information, such as the template structure used, making it difficult to assign functional hierarchies, such as from the SCOP database. Further, template structures are usually annotated and help transfer functions to the modeled protein structure. A structural alignment exercise was performed to assign fold information to *C. acnes* (n=2280), where 2280 proteins were aligned against non-redundant protein structures present in PDB (m=26,775). The program TMalign was used to establish the scan. This resulted in a 6,10,47,000 combination of pairs (2280 × 26775), which is roughly equivalent to 61 million pairs of structural alignments (see Sheet S1 (summary stats)). The top folds (Figure S5(a)) in the modeled C. acnes proteome include TIM beta/alpha-barrel, Rossmann(2×3)oid, Ferredoxin-like, OB-fold, UBA-type 3-helical bundle, etc. The top superfamilies (Figure S5(b)) in the modeled proteome include 'Aldolase', 'Dihydropteroate synthetase-like', and 'Phosphoenolpyruvate/pyruvate domain'.

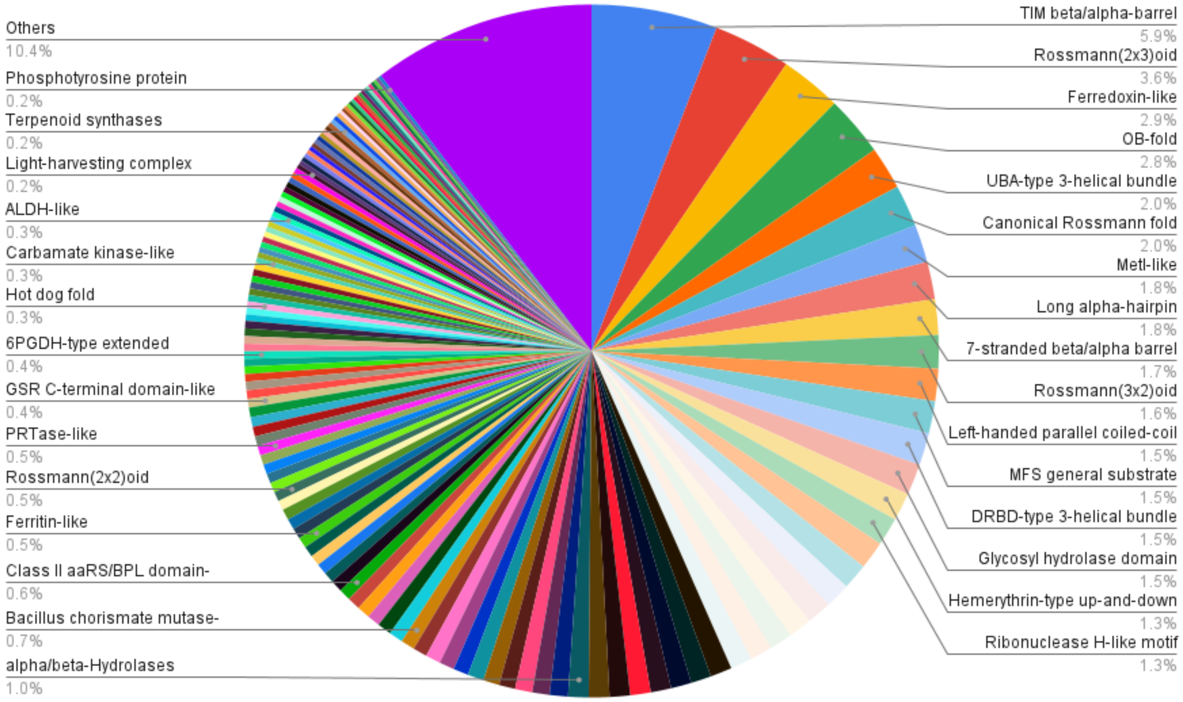


(a) SCOP2: FOLD

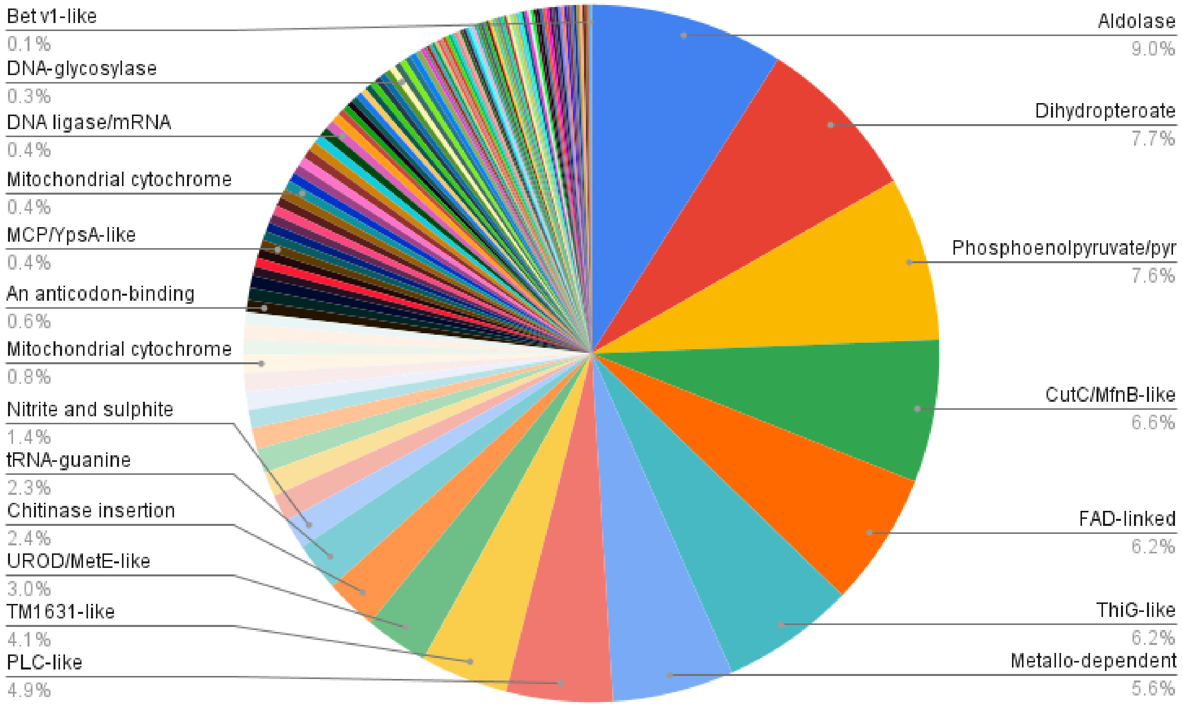


(b) SCOP2: Superfamily

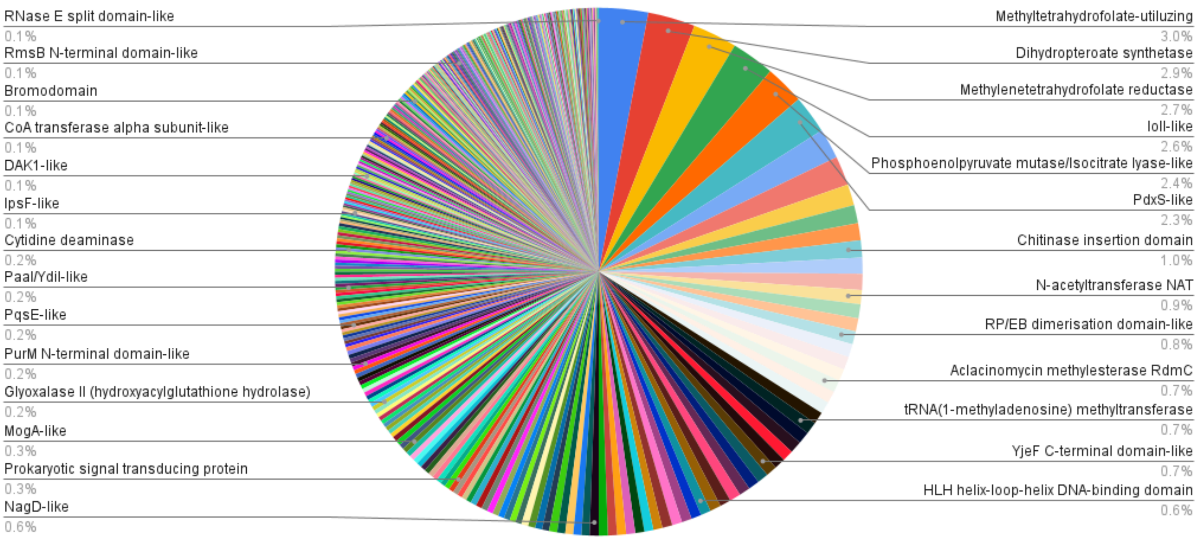


(c) SCOP2: Family

Figure S5: Pie chart displaying the fraction of the modeled proteome associated with (a)Fold, (b)Superfamily, and (c)Family (derived from the SCOP2 database).

| Gene ID | GenBank annotation provided in KEGG database | Annotation in CAcnesDB | Confidence Score | Revised Category | Gene symbol | KEGG pathway ID | KEGG pathway | Functional Class in CAcnesDB | Modeled Structure's Quality | ligand ID | Ligand | Intrermolecular Energy (ΔG) |
|---|---|---|---|---|---|---|---|---|---|---|---|---|
| PPA2105 | (GenBank) triacylglycerol lipase precursor | Triacylglycerol lipase (GehA) | 5 | known | GehA | pac00561, pac01100 | Glycerolipid metabolism, Metabolic pathways | lipid metabolism | 286 (84.37%) | PLM; B12 | PALMITIC ACID; COBALAMIN | PPA2105a_PLM (-7.50); PPA2105a_B12 (Nil) |
| PPA0075 | (GenBank) hypothetical protein | Putative S-adenosyl-L-methionine-dependent (SAM) methyltransferase | 5 | putative | PPA0075 | pac01100 | Metabolic pathways | intermediary metabolism and respiration | 250 (92.59%) | SAM; D3H | S-ADENOSYLMETHIONINE; ~{N}-(4-aminophenyl)-2-azanyl-ethanamide | PPA0075a_D3H (-6.04); PPA0075a_SAM (-7.47) |
| PPA0255 | (GenBank) hypothetical protein | Putative S-adenosyl-L-methionine-dependent (SAM) methyltransferase | 3 | putative | PPA0255 | | | intermediary metabolism and respiration | 307 (88.22%) | SAM; HEM | S-ADENOSYLMETHIONINE; PROTOPORPHYRIN IX CONTAINING FE | PPA0255a_SAM (-8.37); PPA0255a_HEM (-6.43) |
| PPA2025 | (GenBank) hypothetical protein | Putative FMN reductase (NADH) RutF | 4 | putative | PPA2025 | pac01100 | Metabolic pathways | intermediary metabolism and respiration | 162 (72.97%) | FAD; LES; FAD; AO8; BLA | FLAVIN-ADENINE DINUCLEOTIDE; (3S)-N-(pyrimidin-2-yl)azepan-3-amine; FLAVIN-ADENINE DINUCLEOTIDE; [2-[[2-(isoquinolin-5-ylsulfonylamino)ethylamino]methyl]phenyl]boronic acid; BILIVERDINE IX ALPHA | PPA2025a_LES (-4.67); PPA2025a_AO8 (-8.21); PPA2025a_FAD (-3.32); PPA2025a_BLA (-9.42) |
| PPA1036 | (GenBank) hypothetical protein | Putative GNAT family N-acetyltransferase | 4 | putative | PPA1036 | pac01100 | Metabolic pathways | intermediary metabolism and respiration | 71 (86.59%) | COA | COENZYME A | PPA1036a_COA (-7.58) |
| PPA1995 | (GenBank) hypothetical protein | Putative GNAT family N-acetyltransferase | 4 | putative | PPA1995 | pac01100 | Metabolic pathways | intermediary metabolism and respiration | 285 (87.69%) | ACO; FAS; GNP | ACETYL COENZYME *A; ARABINO-FLAVIN-ADENINE DINUCLEOTIDE; PHOSPHOAMINOPHOSPHONIC ACID-GUANYLATE ESTER | PPA1995a_FAS (-4.65); PPA1995a_GNP (-3.94); PPA1995a_ACO (-8.48) |
| PPA2401 | (GenBank) hypothetical protein | Putative GNAT family N-acetyltransferase | 4 | putative | PPA2401 | pac01100 | Metabolic pathways | intermediary metabolism and respiration | 159 (96.36%) | ACO; NAP | ACETYL COENZYME *A; NADP NICOTINAMIDE-ADENINE-DINUCLEOTIDE PHOSPHATE | PPA2401a_ACO (-11.93); PPA2401a_NAP (-5.36) |
| PPA0410 | (GenBank) conserved protein | O-acetyl-ADP-ribose deacetylase | 5 | known | ymdB | pac00230, pac01100 | Purine metabolism, Metabolic pathways | intermediary metabolism and respiration | 170 (76.23%) | APR; ANL; RVP | ADENOSINE-5-DIPHOSPHORIBOSE; ANILINE; RIBAVIRIN MONOPHOSPHATE | PPA0410a_RVP (-4.87); PPA0410a_APR (-9.38); PPA0410a_ANL (-4.44) |
| PPA1407 | (GenBank) biotin synthase | Putative type I Biotin synthase | 6 | known | bioB | pac00780, pac01100, pac01240, pac_M00123, pac_M00577 | Biotin metabolism, Metabolic pathways, Biosynthesis of cofactors, Biotin biosynthesis, pimeloyl-ACP/CoA => biotin, Biotin biosynthesis, BioW pathway, pimelate => pimeloyl-CoA => biotin | intermediary metabolism and respiration | 326 (84.02%) | SF4; SAH; NAD | IRON/SULFUR CLUSTER; S-ADENOSYL-L-HOMOCYSTEINE; NICOTINAMIDE-ADENINE-DINUCLEOTIDE | PPA1407a_SAH (-4.31); PPA1407a_NAD (-6.15); PPA1407a_SF4 (-3.57) |
| PPA0731 | (GenBank) hypothetical protein | Putative barstar family protein | 3 | putative | PPA0731 | pac00910, pac01100 | Nitrogen metabolism, Metabolic pathways | regulatory protein | 103 (78.03%) | FAD; 1O0 | FLAVIN-ADENINE DINUCLEOTIDE; 6-[3-(pyridin-2-yl)-2-(trifluoromethyl)phenoxy]-1-(2,4,6-trifluorobenzyl)quinazolin-4(1H)-one | PPA0731a_1O0 (-6.28); PPA0731a_FAD (-4.09) |
| PPA2334 | (GenBank) hypothetical protein | Ppi-type phosphoenolpyruvate carboxykinase (diphosphate) | 3 | putative | PPA2334 | pac00620, pac01100 | Pyruvate metabolism, Metabolic pathways | intermediary metabolism and respiration | 1108 (97.79%) | U0E; ACP; A82; B3I | N-[[1-[N-ACETAMIDYL]-[1-CYCLOHEXYLMETHYL-2-HYDROXY-4-ISOPROPYL]-BUT-4-YL]-CARBONYL]-GLUTAMINYL-ARGINYL-AMIDE; PHOSPHOMETHYLPHOSPHONIC ACID ADENYLATE ESTER; 2-[[(1R)-1-(7-methyl-2-morpholin-4-yl-4-oxidanylidene-pyrido[1,2-a]pyrimidin-9-yl)ethyl]amino]benzoic acid; 2,3,5-TRIIODOBENZOIC ACID | PPA2334a_ACP (-6.32); PPA2334a_A82 (+43.55); PPA2334a_B3I (-7.64); PPA2334a_U0E (-11.62) |
| PPA2204 | (GenBank) hypothetical protein | Transcription factor WhiB | 3 | putative | PPA2204 | pac02020 | Two-component system | regulatory protein | 152 (82.16%) | ATP; FAD | ADENOSINE-5'-TRIPHOSPHATE; FLAVIN-ADENINE DINUCLEOTIDE | PPA2204a_FAD (-6.15); PPA2204a_ATP (-4.32) |
| PPA2023 | (GenBank) hypothetical protein | Putative Phosphomannomutase from HAD superfamily | 3 | putative | PPA2023 | pac00051, pac00520, pac01100, pac01110 | Fructose and mannose metabolism, Amino sugar and nucleotide sugar metabolism, Metabolic pathways, Biosynthesis of secondary metabolites | intermediary metabolism and respiration | 223 (99.55%) | CIT; PZH | CITRIC ACID; 1-(4-BROMOPHENYL)METHANAMINE | PPA2023a_PZH (-5.74); PPA2023a_CIT (-2.73) |
| PPA0047 | (GenBank) hypothetical membrane-spanning protein | Putative LysM peptidoglycan-binding domain-containing protein | 3 | putative | PPA0047 | pac00230, pac00240, pac01100 | Purine metabolism, Pyrimidine metabolism, Metabolic pathways | Cell wall and cell processes | 636 (69.13%) | 3CL; 5RC; NAI | 3-CHLOROPROPANOL; ~{N}-[(1~{R},6~{R})-6-azanyl-2,2-bis(fluoranyl)cyclohexyl]-5-ethyl-4-[6-(trifluoromethyl)pyrazolo[1,5-a]pyrimidin-3-yl]thiophene-2-carboxamide; 1,4-DIHYDRONICOTINAMIDE ADENINE DINUCLEOTIDE | PPA0047a_5RC (-10.54); PPA0047a_NAI (-4.81); PPA0047a_3CL (-3.17) |
| PPA0231 | (GenBank) hypothetical protein | Putative pilus assembly protein TadB | 4 | putative | PPA0231 | pac02020 | Two-component system | Cell wall and cell processes | 204 (79.69%) | BB2; CLR; ARC; HEM | ACTINONIN; CHOLESTEROL; 3,7,11,15-TETRAMETHYL-HEXADECAN-1-OL; PROTOPORPHYRIN IX CONTAINING FE | PPA0231a_BB2 (-6.29); PPA0231a_CLR (-8.00); PPA0231a_ARC (-5.17); PPA0231a_HEM (-5.09) |
| PPA0064 | (GenBank) putative hydrolase | Putative Fructose-1-phosphate phosphatase YqaB, HAD family phosphatase | 5 | putative | PPA0064 | pac00740, pac01100, pac01110, pac01240 | Riboflavin metabolism, Metabolic pathways, Biosynthesis of secondary metabolites, Biosynthesis of cofactors | intermediary metabolism and respiration | 207 (96.73%) | PGA | 2-PHOSPHOGLYCOLIC ACID | PPA0064a_PGA (-1.84) |
| PPA0768 | (GenBank) pseudouridylate synthase | Pseudouridine synthase (RsuA/RluA) | 7 | known | PPA0768 | pac00740, pac01100 | Riboflavin metabolism, Metabolic pathways | information pathways | 295 (96.41%) | U; 6UL; ADP; FAD | URIDINE-5'-MONOPHOSPHATE; TETRACOSYL PALMITATE; ADENOSINE-5'-DIPHOSPHATE; FLAVIN-ADENINE DINUCLEOTIDE | PPA0768a_FAD (-8.55); PPA0768a_U (-8.48); PPA0768a_ADP (-5.13); PPA0768a_6UL (-4.42) |

Table S2: The table lists the examples with consolidated annotations from both sequence- and structure-based analyses.

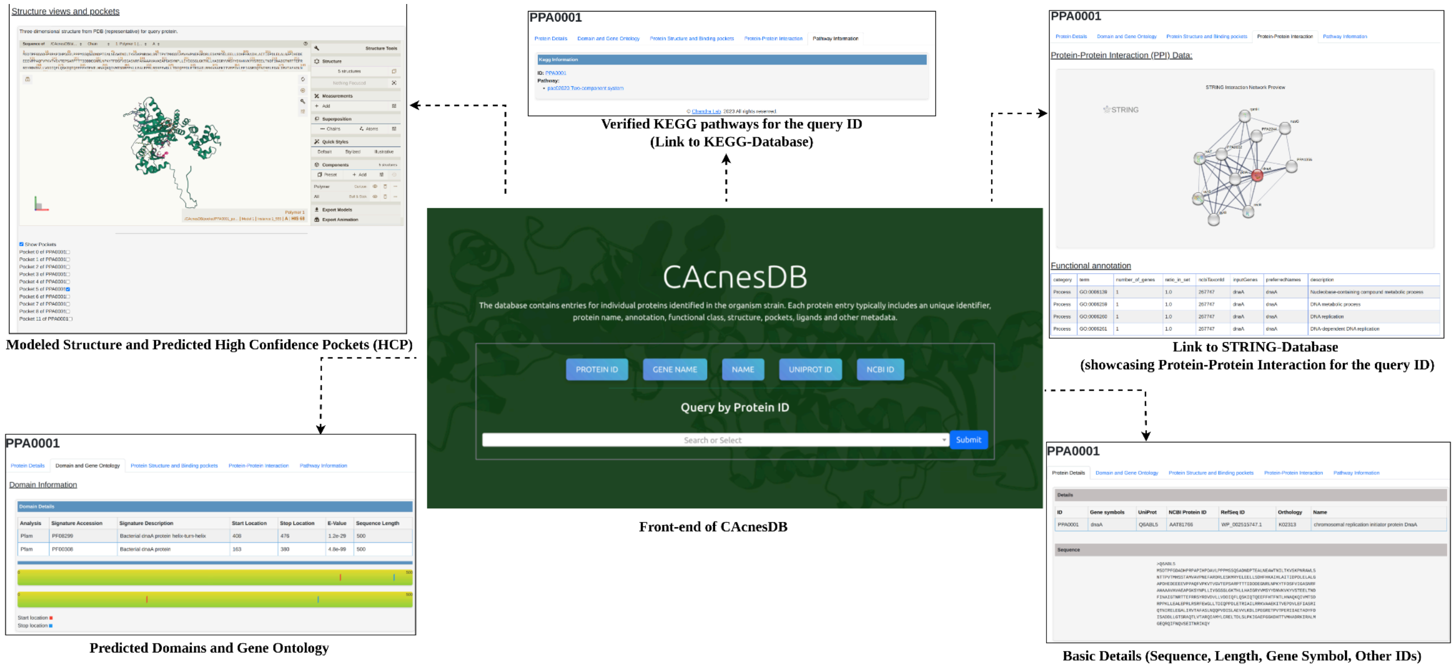


Figure S6: Front-end of CAcnesDB showcasing different functionalities.
